# Naming-related spectral responses predict neuropsychological outcome after epilepsy surgery

**DOI:** 10.1101/2021.04.11.439389

**Authors:** Masaki Sonoda, Robert Rothermel, Alanna Carlson, Jeong-Won Jeong, Min-Hee Lee, Takahiro Hayashi, Aimee F. Luat, Sandeep Sood, Eishi Asano

**Affiliations:** Department of Pediatrics, Children’s Hospital of Michigan, Detroit Medical Center, Wayne State University, Detroit, Michigan, 48201, USA; Department of Neurology, Children’s Hospital of Michigan, Detroit Medical Center, Wayne State University, Detroit, Michigan, 48201, USA; Department of Psychiatry, Children’s Hospital of Michigan, Detroit Medical Center, Wayne State University, Detroit, Michigan, 48201, USA; Department of Neurosurgery, Children’s Hospital of Michigan, Detroit Medical Center, Wayne State University, Detroit, Michigan, 48201, USA; Department of Neurosurgery, Yokohama City University, Yokohama, Kanagawa, 2360004, Japan

**Keywords:** pediatric epilepsy surgery, intracranial electroencephalography (iEEG) recording, event-related high gamma augmentation, high-frequency oscillations (HFOs), ripples

## Abstract

This prospective study determined the utility of intracranially-recorded spectral responses during naming tasks in predicting neuropsychological performance following epilepsy surgery. We recruited 65 patients with drug-resistant focal epilepsy who underwent preoperative neuropsychological assessment and intracranial EEG (iEEG) recording. The Clinical Evaluation of Language Fundamentals (CELF) evaluated the baseline and postoperative language function. During extraoperative iEEG recording, we assigned patients to undergo auditory and picture naming tasks. Time-frequency analysis determined the spatiotemporal characteristics of naming-related amplitude modulations, including high gamma augmentation (HGA) at 70-110 Hz. We surgically removed the presumed epileptogenic zone based on the extent of iEEG and MRI abnormalities while maximally preserving the eloquent areas defined by electrical stimulation mapping (ESM). The multivariate regression model incorporating auditory naming-related HGA predicted the postoperative changes in Core Language Score (CLS) on CELF with r^2^ of 0.37 (p = 0.015) and in Expressive Language Index (ELI) with r^2^ of 0.32 (p = 0.047). Independently of the effects of epilepsy and neuroimaging profiles, higher HGA at the resected language-dominant hemispheric area predicted a more severe postoperative decline in CLS (p = 0.004) and ELI (p = 0.012). Conversely, the model incorporating picture naming-related HGA predicted the change in Receptive Language Index (RLI) with r^2^ of 0.50 (p < 0.001). Higher HGA independently predicted a more severe postoperative decline in RLI (p = 0.03). Ancillary regression analysis indicated that naming-related low gamma augmentation as well as alpha/beta attenuation likewise independently predicted a more severe CLS decline. The machine learning-based prediction model, referred to as the boosted tree ensemble model, suggested that naming-related HGA, among all spectral responses utilized as predictors, most strongly contributed to the improved prediction of patients showing a >5-point CLS decline (reflecting the lower 25 percentile among patients). We generated the model-based atlas visualizing sites, which, if resected, would lead to such a CLS decline. The auditory naming-based model predicted patients who developed the CLS decline with an accuracy of 0.80. The model indicated that *virtual resection* of an ESM-defined language site would have increased the relative risk of the CLS decline by 5.28 (95%CI: 3.47 to 8.02). Especially, that of an ESM-defined receptive language site would have maximized it to 15.90 (95%CI: 9.59-26.33). In summary, naming-related spectral responses predict objectively-measured neuropsychological outcome after epilepsy surgery. We have provided our prediction model as an open-source material, which will indicate the postoperative language function of future patients and facilitate external validation at tertiary epilepsy centers.

## INTRODUCTION

Invasive recording using intracranial electrodes aims to localize the seizure onset zone (SOZ) and functionally-important areas for resective epilepsy surgery.^1^ Electrical stimulation mapping (ESM) remains the gold standard for defining the extent of language areas.^2–5^ Investigators have implemented an alternative method for language mapping partly because ESM has several limitations. Stimulation of nonepileptic areas may induce non-habitual seizures,^6^ which may increase the risk of surgical complications and reduce the reliability of the subsequent ESM sessions. A previous study of 122 patients reported that ESM induced seizures in 36% of patients.^7^ Some patients may not be able to tolerate ESM sessions lasting hours. A tertiary epilepsy center reported that ESM failed to localize the language areas in the presumed dominant hemisphere in four-fifths of children at the age of 10 years or younger.^8^

Measurement of language task-related spectral responses on intracranial EEG (iEEG) is a method that complements the gold-standard ESM.^9–12^ Task-related augmentation of high-frequency broadband activity, including high gamma (70-110 Hz), is considered to reflect cortical activation at a given moment.^13,14^ Such amplitude augmentation was reported to be associated with increased local neural firing^15,16^ hemodynamic activation on functional MRI,^17^ and increased metabolism on glucose positron emission tomography.^18^ A meta-analysis of 15 studies reported that electrode sites showing naming-related high gamma augmentation had 6.44 times increased odds to be classified as the ESM-defined language areas.^19^ Our preliminary study reported that resectioning sites showing naming-related high gamma augmentation increased the risk of new language deficits requiring speech therapy, independent of objective neuropsychological assessment.^20^ Two iEEG studies of 11 and 17 patients recently reported that resection of high gamma sites was marginally associated with a postoperative decline of neuropsychological function.^21,22^ A prospective study of a larger cohort of patients with neuropsychological data is necessary to provide definitive evidence supporting the clinical utility of high gamma-based language mapping.

The present study intended to achieve the following three aims. **[Aim 1]** We aimed to clarify the causal relationship between cortical resection involving naming-related high gamma activation sites and *<u>objectively-measured neuropsychological performance</u>* following epilepsy surgery. We specifically determined whether high gamma-based mapping would predict postoperative language performance *<u>independently of</u>* epilepsy and neuroimaging profiles available preoperatively. **[Aim 2]** We determined whether naming-related modulations of iEEG frequency bands other than high gamma would likewise predict postoperative language performance. Previous iEEG studies reported that sites showing high gamma augmentation frequently also exhibit augmentation of low gamma activity as well as suppression of alpha/beta activity.^23–25^ While high gamma augmentation is suggested to be better time-locked to a given task than alpha/beta suppression,^13,26^ it remains to be determined whether high gamma-based mapping would play the most critical role in predicting postoperative language function. **[Aim 3]** We generated a machine learning-based prediction model identifying electrode sites, which, if resected, would lead to a postoperative decline in language function. We internally validated our prediction model by demonstrating concordance with the ESM findings. Specifically, our simulation-based assessment determined the relative risk of a postoperative language decline related to *<u>virtual</u>* resection of an ESM-defined language site. To facilitate external validation, we have provided our prediction model as open-source material. Investigators can use this model to predict a given patient’s postoperative language outcome at their own epilepsy centers.

## MATERIALS AND METHODS

### Patients

We prospectively recruited and studied a consecutive series of patients satisfying the following criteria. The inclusion criteria consisted of [a] drug-resistant focal epilepsy, [b] age four years and above, [c] neuropsychological evaluation including the baseline language function,^27^ [d] extraoperative iEEG recording as part of our presurgical evaluation at Detroit Medical Center in Detroit between January 2009 and February 2019, and [e] measurement of naming-related spectral responses on iEEG. The exclusion criteria consisted of [a] history of previous epilepsy surgery and [b] massive structural lesions (such as megalencephaly or perisylvian polymicrogyria), which would make the Sylvian or central sulcus unidentifiable. The Wayne State University Institutional Review Board approved the study. We obtained informed consent/assent in writing from the patients or the guardians of patients.

### Language dominant hemisphere

We previously discussed the rationale of our approach to estimate the language dominant hemisphere in children.^20,28,29^ It is infeasible to expect all surgical candidates, especially young children, would successfully undergo functional MRI-based lateralization of the language dominant hemisphere. Thus, we estimated the dominant hemisphere based on the handedness and anatomical MRI findings.^30–32^ We treated the left hemisphere as dominant if the patient was [1] right-handed or [2] left-handed but free of a developmental cortical lesion (such as dysplasia) in the left neocortical area. Conversely, we treated the right hemisphere as dominant if the patient was left-handed and had a developmental cortical lesion in the left neocortex. The present study aimed to generate and validate a model that can predict postoperative language function without relying on functional MRI studies or ESM.^8,33^

### iEEG

We surgically implanted platinum disk electrodes on the pial surface to determine the boundary between the SOZ and functionally-important areas.^28,34^ We continuously recorded iEEG at the bedside with a sampling rate of 1,000 Hz and a band-pass of 0.016-300 Hz for 3-7 days.^35^ We discontinued antiepileptic drugs (AEDs) to capture habitual spells and localize the SOZ responsible for generating habitual seizures.^34^ We performed the following iEEG analysis using common average reference (i.e., an average of iEEG voltages at all channels excluding those affected by SOZ, interictal spikes, MRI lesions, or artifacts).

### MRI

Before implanting intracranial electrodes, we acquired 3T MRI, including a T1-weighted spoiled gradient-echo volumetric scan and fluid-attenuated inversion recovery scan.^36^ We coregistered electrodes with a three-dimensional surface image^37,38^ Furthermore, we spatially normalized all electrode locations of all patients to the FreeSurfer averaged image (http://surfer.nmr.mgh.harvard.edu).^35,37,39^ **Figure 1** shows the spatial distribution of intracranial electrodes included in our iEEG analysis.

**Figure 1.**
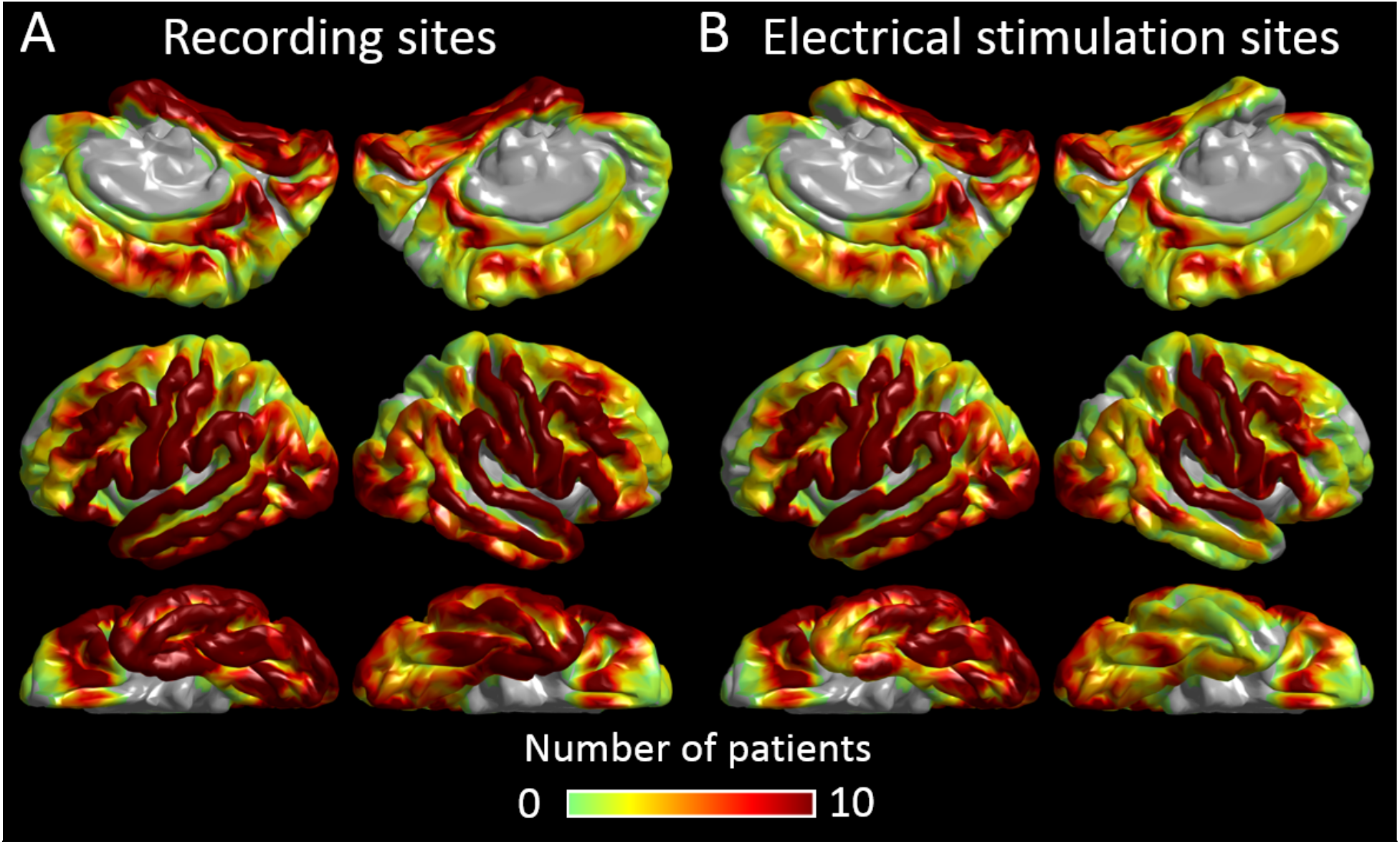
Distribution of intracranial electrodes. **A,** The FreeSurfer surface image presents the distribution of artifact-free electrode sites included in the present study (6,886 sites). Color indicates the number of patients at each cortical point. **B,** The distribution of sites assessed by electrical stimulation mapping (5,203 sites).

### ESM

We previously described our ESM protocol in detail.^6,37^ We stimulated a pair of neighboring electrode sites with a frequency of 50 Hz, pulse width of 0.3 ms, train duration of ≤5 s. **Figure 1** shows the distribution of 5,203 electrode sites assessed by ESM. We initially set the stimulus intensity at 3 mA, increased it to 6 and 9 mA in a stepwise manner until a clinical symptom or afterdischarge was noted. We kept the stimulus intensity below the afterdischarge threshold once identified in a given patient. During each stimulation trial, a given patient was instructed to answer auditory questions such as ‘*What flies in the sky?*’ or to name pictures presented by a neuropsychologist (R.R.) blinded to the results of naming-related spectral responses. The neuropsychologist asked each patient what made her/him fail to respond when needed. Additional tasks (e.g., syllable repetition) specified the functional role of each stimulated site. Sites at which stimulation induced the following symptoms were defined as ESM-defined language areas. [a] Speech arrest: inability to vocalize. [b] Auditory receptive aphasia: failure to understand auditory questions. [c] Auditory expressive aphasia: intact vocalization, successful understanding of auditory questions, but failure to provide a relevant answer. [d] Visual expressive aphasia: intact vocalization but failure to name pictures. We visualized the group-level probability of stimulation-induced symptoms at each cortical point on the FreeSurfer averaged surface image.^37^

### Surgery

All patients underwent epilepsy surgery within 24 hours after the completion of extraoperative iEEG recording. Our primary intention was to completely remove the presumed epileptogenic zone consisting of SOZ and the neighboring structural lesion while maximally preserving the eloquent areas.^34,36,40^ The spatial extent of ESM-defined language areas and the spatiotemporal profiles of naming-related high-gamma augmentation were available before the surgery. We considered that ESM would localize the regions *<u>essential</u>* for language, whereas naming-related high-gamma augmentation would localize those *<u>involved</u>* in language.^20,28^ With the patient and family, before the intracranial electrode placement as well as at least a day before the surgery, we discussed the pros and cons of the complete and incomplete resection of the presumed epileptogenic zone in case the language areas were spatially overlapped with the SOZ.^34^

Based on the intraoperative photograph taken immediately before the dural closure, the FreeSurfer script computed the resection size, defined as the proportion of resected tissue among the hemisphere (% of the hemisphere).^36^ We previously reported that the resection size estimated with an intraoperative photograph was highly concordant with that based on postoperative MRI.^40^

### Neuropsychological assessment

A neuropsychologist (A.C.), being blinded to the results of any iEEG analysis, evaluated the preoperative and postoperative language function using the Clinical Evaluation of Language Fundamentals-Fourth Edition (CELF-4).^27^ We computed the age-corrected Core Language Score (CLS), Receptive Language Index (RLI), and Expressive Language Index (ELI) (average: 100; standard deviation: 15) for patients with ages between 4 and 21 years. Since a given patient was expected to gradually recover from postoperative language impairment, if any, as a function of time, we treated the interval between surgery and postoperative neuropsychological assessment (**Table 1**) as a covariate in the multivariate regression analysis below.

**Table 1.**
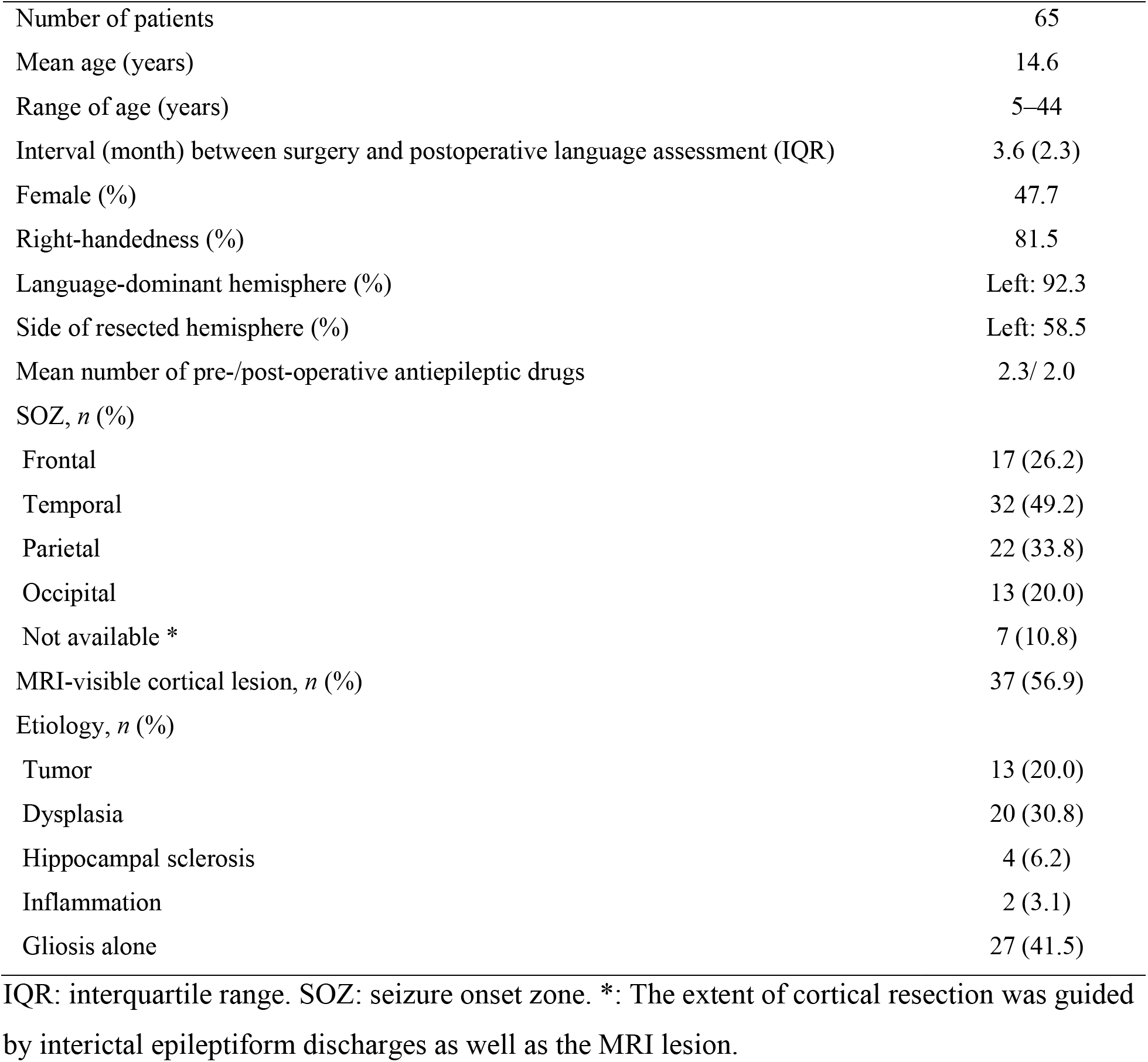
Patient profile.

### Naming tasks

We assigned auditory and picture naming tasks to patients during interictal iEEG recording to localize the cortical sites *<u>involved</u>* in language based on the time-frequency analysis.^28,29^ Our previous studies described the task parameters in detail.^29,37^ We synchronized iEEG traces, stimulus presentations, and patient behaviors using a photosensor and microphones.^28^ For the auditory naming task, we instructed patients to overtly provide an answer for each of up to 100 audible sentence questions. We measured the percentage of correct answers and the response time defined as the interval between stimulus offset and response onset (**Fig. 2**). We excluded trials from time-frequency analysis if a given patient failed to provide a relevant answer or the response time was longer than two standard deviations from the individual mean.^41^

**Figure 2.**
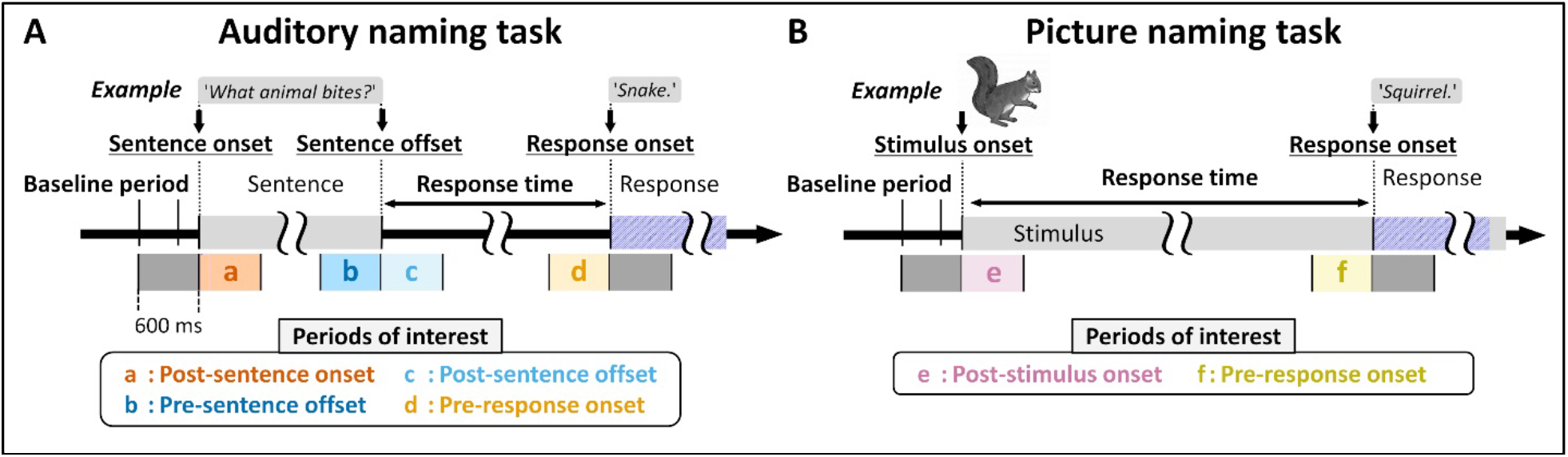
Naming tasks and analysis periods of interest. **A,** Auditory naming task. Each patient was instructed to verbally answer a brief sentence question (median duration of sentence stimuli: 1.8 s; range: 1.2 to 2.4 s). **B,** Picture naming task. Each patient named an object presented on a monitor. Each 600-ms analysis period of interest is highlighted in color and labeled as **a** to **f**.

For the picture naming task, we asked patients to overtly name an object presented on an LCD monitor (up to 60 common objects such as ‘dog’ and ‘tree’). We likewise measured the percentage of correct answers and the response time defined as the interval between stimulus onset and response onset. We aligned iEEG traces to stimulus onset and response onset (**Fig. 2**).

### Measurement of naming-related high gamma responses

At each artifact-free channel, we determined the temporal profiles of iEEG high gamma (70-110Hz) amplitude augmentation during the auditory naming task. We measured amplitude modulations during 1,200-ms epochs centered at sentence onset, sentence offset, and response onset, whereas picture naming-related responses during 1,200-ms epochs centered at stimulus onset and response onset (**Fig. 2**). Using the Morlet wavelet method implemented in FieldTrip (http://www.fieldtriptoolbox.org/), we transformed iEEG voltage data into time-frequency bins (1 Hz frequency bins ranging from 2 to 110 Hz; each frequency × 0.1 cycles) sliding in 10 ms steps.

We minimized the unwanted direct effect of interictal spikes on naming-related high gamma modulations by removing the time-frequency bins showing an excessive and irregular increase in broadband amplitude. If the amplitude averaged across 30-85 Hz at a given time in a trial was greater than two standard deviations from the mean amplitude across all trials in a given patient, we treated the whole time-frequency bins at the corresponding time on that trial as missing values.^42,43^ We believe this analytic approach can effectively extract and eliminate the pathological high gamma component carried by interictal spike discharges, which would randomly occur *<u>without</u>* being time-locked to stimuli or responses (**Fig. 3**).

**Figure 3.**
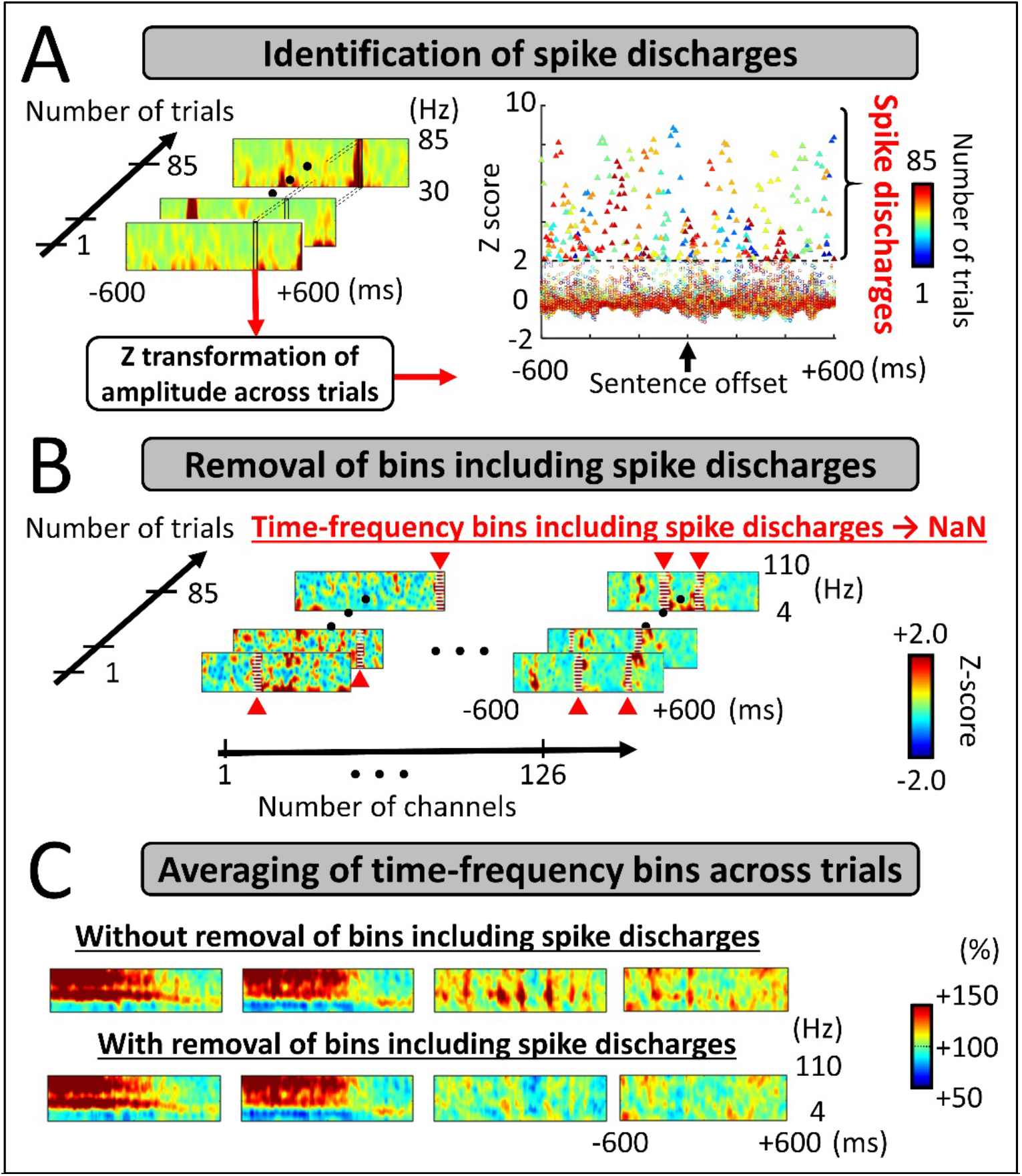
Preprocessing to remove pathological components from time-frequency analysis. **A,** Identification of randomly occurring spike discharges. We identified within-trial time-frequency bins showing a broadband (30-85 Hz) amplitude greater than two standard deviations from the across-trial mean (i.e., z-score of >2).^42,43^ **B,** Removal of bins including spike discharges. We treated such timefrequency bins including an excessive broadband amplitude as missing values (i.e., NaN). **C,** Averaging of time-frequency bins across trials. We computed the averaged amplitude modulations (i.e., percent change) as compared to that during the 400-ms baseline period prior to the stimulus onset.^37^ Here, the time-frequency matrices present auditory naming-related spectral responses time-locked to stimulus offset at four electrode sites. Upper: Across-trial averaged data before bins including spike discharges excluded; several matrices exhibit episodes of brief broadband augmentation attributed to randomly-occurring spike discharges. Lower: Across-trial averaged data after excluding bins with spike discharges. In the present study, we adopted the time-frequency data presented in the lower row. Amplitude scale: 100% indicates no change in amplitude compared to the baseline, whereas 110% and 90% indicate 10% increase and decrease, respectively.

We subsequently determined when naming-related high gamma augmentation reached significance at each electrode site. We tested the null hypothesis that high gamma amplitude at each 10-ms bin would be the same as that during the baseline period with a two-sided 5% significance level (permutation test [n = 500] with FDR correction for repeated comparisons for 121 bins in a 1,200-ms period; **Fig. 2**). We treated bins showing amplitude augmentation for at least three consecutive high gamma cycles (i.e., >33 ms) as significant high gamma augmentation. We finally computed auditory naming-related high gamma augmentation averaged across the four analysis periods of interest (**Fig. 2A**) at sites showing significant high gamma augmentation. Likewise, we computed picture naming-related high gamma augmentation averaged across the two analysis periods of interest (**Fig. 2B**). These high gamma values were treated as the summary measure reflecting the degree of local task-related cortical activation and incorporated in the subsequent multivariate regression analysis.

### Statistical analysis: iEEG high gamma and ESM

We determined whether the temporal profile of naming-related high gamma augmentation would account for language symptoms elicited by ESM. Using the Spearman’s rank correlation coefficient, we visualized how well the degree of significant high gamma augmentation during a specific 600-ms analysis period relative to stimulus and response would be correlated to the probability of each of the ESM-induced language symptoms (**Fig. 4**).

**Figure 4.**
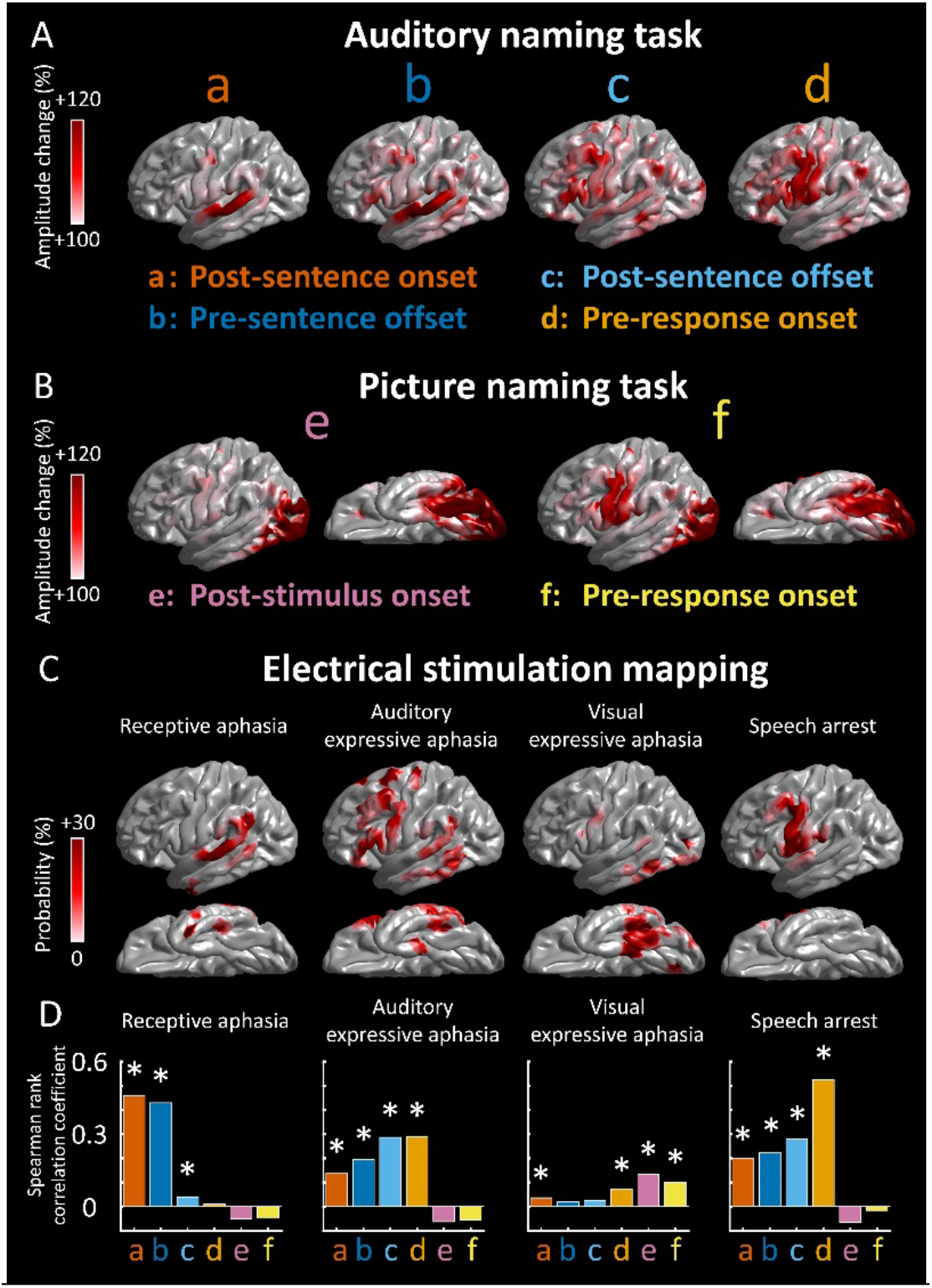
Naming-related high gamma augmentation and electrical stimulation mapping (ESM). **A**, The spatial distribution of auditory naming-related high gamma augmentation during each 600-ms analysis period of interest: **a**, post-sentence onset; **b**, pre-sentence offset; **c**, post-sentence offset; **d**, pre-response onset. **B**, The distribution of picture naming-related high gamma augmentation: **e**, post-stimulus onset; **f**, pre-response onset. **C**, The spatial distribution of the probability of ESM-induced symptoms, including receptive aphasia, auditory expressive aphasia, visual expressive aphasia, and speech arrest. **D,** The bar charts show the strength of correlation between modality-specific high gamma augmentation (at sites with a z-score of ≥2) and the probability of each ESM-induced symptom. *: Positive correlation with a Bonferroni-adjusted p <

#### Statistical analysis: iEEG high gamma and neuropsychological data

**[Aim 1]** Multivariate linear regression analysis determined whether high gamma-based mapping would predict postoperative language performance *<u>independently of</u>* epilepsy and neuroimaging data available preoperatively.^44^ We used MATLAB 2020a Statistics and Machine Learning Toolbox (MathWorks, Natick, MA, USA) and set significance at p<0.05. The predictor variables included: [#1] ‘maximum resected high gamma (%)’ defined as the high gamma percent change highest among sites included in the resected language-dominant hemispheric region, [#2] the resection size of the language-dominant hemispheric cortex (%), [#3] age at surgery (years), [#4] sex (1 if female), [#5] interval between surgery and postoperative neuropsychological assessment (months), [#6] number of oral antiepileptic drugs taken preoperatively (reflecting the severity of epilepsy-burden),^40,45^ [#7] MRI-visible cortical lesion (1 if present), [#8] SOZ location (1 if frontal or temporal), and [#9] preoperative CELF-4 score. With a sample size of 65, a power of 0.8, an alpha of 0.05, and nine predictors incorporated, the regression model was anticipated to detect a moderate effect size of *f*^2^ of 0.28. We assumed that the test-retest reproducibility would be comparable across patients.^46,47^

### Statistical analysis: Utility of iEEG low gamma, beta, and alpha modulations

**[Aim 2]** As an ancillary analysis, we determined whether task-related modulations of iEEG frequency bands lower than high gamma would likewise predict postoperative language performance. Using the aforementioned time-frequency analysis to measure naming-related high gamma augmentation, we computed low gamma augmentation (30-50 Hz), beta attenuation (12-30 Hz), and alpha attenuation (8-12 Hz) during a given naming task. The multivariate linear regression analysis likewise determined whether ‘maximum resected low gamma’, ‘maximum resected beta’, or ‘maximum resected alpha’ would predict postoperative language performance independently of the other covariates mentioned above.

### Machine learning: iEEG amplitude modulations and neuropsychological data

**[Aim 3]** We generated the machine learning-based atlas visualizing the sites, which, if resected, would lead to a postoperative decline in CELF-4-based language function. For this purpose, we used the ensemble learning algorithm supported by Statistics and Machine Learning Toolbox implemented in the MATLAB R2020a (https://www.mathworks.com/help/stats/ensemble-algorithms.html). To accurately predict patients with a postoperative decline in CLS by more than five points (i.e., reflecting the lower 25 percentile in our study cohort), we generated the boosted tree ensemble model initially based on the auditory naming-related amplitude modulations. The model incorporated the following 17 predictors measured during the auditory naming task: [#1 to #4] ‘maximum resected high gamma augmentation (%)’, here defined as the high gamma percent change, in each of the four analysis periods (**Fig. 2A**), highest among electrode sites within the resected region, [#5 to #8] ‘maximum resected low gamma augmentation (%)’, [#9 to #12] ‘maximum resected beta attenuation (%)’, [#13 to #16] ‘maximum resected alpha attenuation (%)’, and [#17] language dominance of the resected hemisphere (i1 if resection involved the dominant hemisphere). Thereby, we linearly zero-centered auditory naming-related amplitude modulations (i.e., 0% reflects no augmentation or attenuation compared to the baseline.^48^ We selected the Gentle Adaptive Boosting as the ensemble learning algorithm.^49^ We subsequently utilized the Bayesian optimization algorithm with the expected-improvement acquisition function, which automatically selected the best set of values of the following hyperparameters through 100 iterations: ‘maximum number of splits’, ‘number of learners’, and ‘number of predictors to sample’. The following article describes the algorithm outline of the Bayesian optimization used in this study (https://www.mathworks.com/help/stats/bayesian-optimization-algorithm.html). We evaluated the prediction performance of our boosted tree ensemble model in predicting patients who developed such a postoperative CLS decline, using [a] accuracy and [b] area under the receiver operating characteristic (ROC) curve in five-fold cross-validation.

Using the MATLAB ‘predictorImportance’ function (https://www.mathworks.com/help/stats/compactregressionensemble.predictorimportance.html),^50^ we determined the relative importance of each variable in predicting patients developing a >5-point CLS decline (**Fig. 5C**).

**Figure 5.**
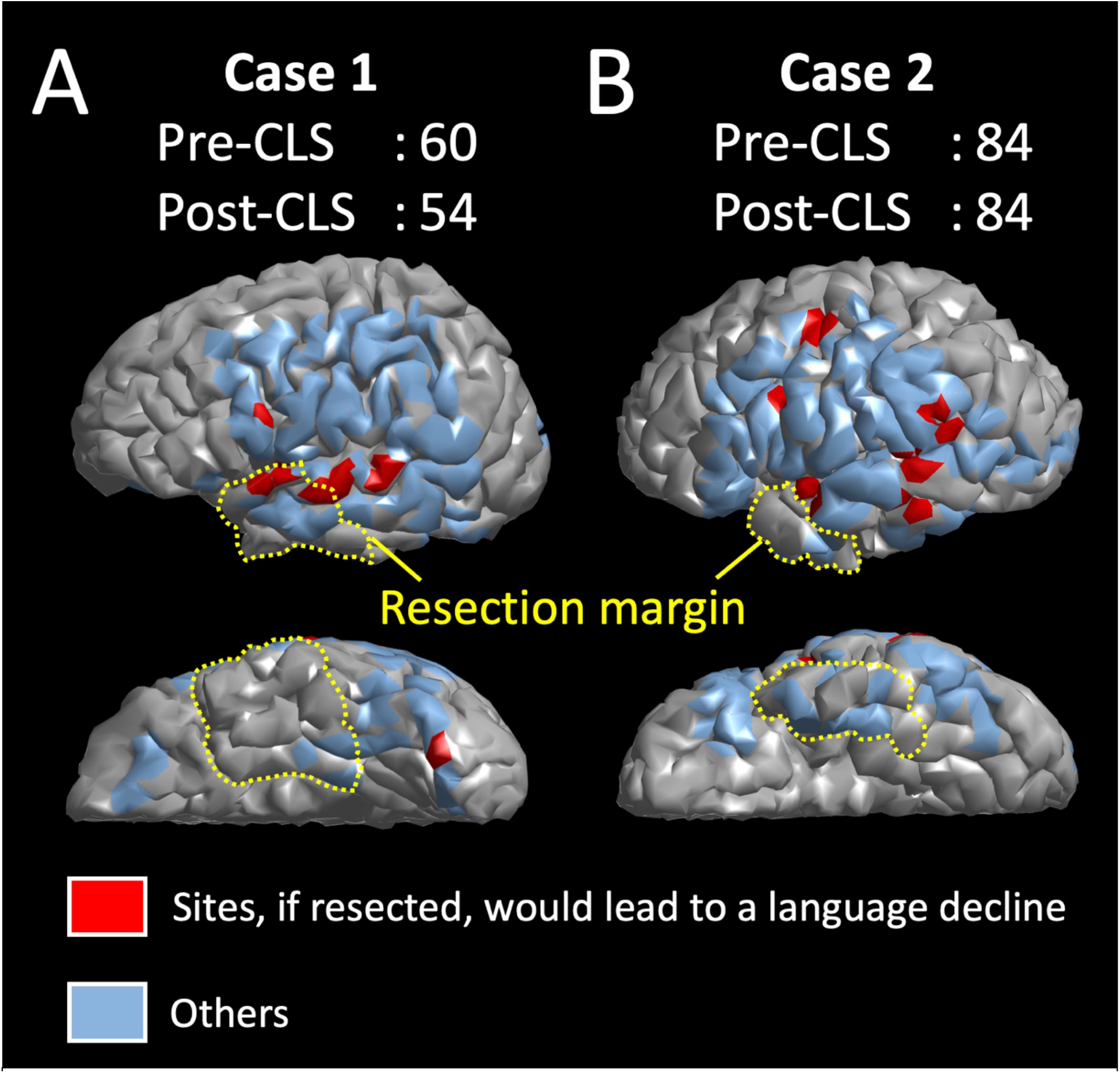
Spatial relationship between cortical prediction sites of postoperative cognitive decline and resection margin. The cortical surface in each individual 3D brain image is highlighted whether a given site, if resected, would result in a >5-point CLS decline. Postoperative cognitive outcome of each electrode site was predicted by the boosted-tree-ensemble model incorporating auditory naming-related iEEG amplitude variables in addition to the dominant hemisphere variable. The yellow dashed lines denote the resection margin in a given patient. **A,** Case 1: The cortical sites predicted to result in a >5-point CLS decline (colored in red) were indeed resected, and the language decline was observed postoperatively. **B,** Case 2: The cortical sites predicted to result in >5 points of CLS decline were preserved, and no language decline was observed.

### Validation of our machine learning-based model using virtual resection of ESM-defined language areas

Our boosted tree ensemble model predicted whether *<u>virtual resection</u>* of a given electrode site would result in a >5-point CLS decline in all 65 study patients. The model was designed to predict the clinical consequence associated with resection of sites in which naming-related spectral responses ranged strictly within those observed in the training data (i.e., a procedure referred to as interpolation.^51^ On the FreeSurfer averaged surface image, we have provided a group-level atlas that visualizes the probability of a >5-point CLS decline resulting from virtual resection of given cortical points (**Fig. 5A**). To validate our boosted tree ensemble model, we computed the relative risk of language impairment resulting from virtual resection of ESM-defined language sites compared to that of the others.^52^ It is feasible to hypothesize that a given patient would develop a substantial language impairment if an ESM-defined language site were surgically removed instead of being preserved.

We likewise generated the boosted tree ensemble model incorporating picture naming-related amplitude modulations. We assessed the model performance in predicting patients who developed a postoperative>5-point CLS decline (**Figs 5B and 5D**). The picture naming-based model incorporated the following nine predictors: [#1 to #2] ‘maximum resected high gamma augmentation (%)’, in each of the two analysis periods (**Fig. 2B**), highest among electrode sites within the resected region, [#3 to #4] ‘maximum resected low gamma augmentation (%)’, [#5 to #6] ‘maximum resected beta attenuation (%)’, [#7 to #8] ‘maximum resected alpha attenuation (%)’, and [#9] language dominance of the resected hemisphere.

### Data and code availability

*<u>All data and code are available upon request to the corresponding author (E.A.).</u>* We are pleased to re-analyze the data based on reviewers’ and readers’ specific suggestions to improve the language mapping method.

## RESULTS

### Patient profiles

A total of 65 patients satisfied the inclusion and exclusion criteria (6,886 artifact-free electrode sites in total; 105.9 per patient on average [SD: ±17.9]). Due to the time constraint during the extraoperative iEEG recording, a single patient failed to complete the auditory naming task and six the picture naming task. Thus, 64 (105.7 ± 17.9) and 59 patients (104.2 ± 22.2) contributed to the analysis of auditory and picture naming-related amplitude modulations (**Fig. 1A**). A total of 5,203 artifact-free electrode sites were assessed by ESM (**Fig. 1B**).

Resective surgery involved a total of 1,938 electrode sites (mean: 29.8 sites per patient; SD: ±24.5). Of the resected electrodes, 1,798 electrodes (92.8%) implanted on the brain surface without structural abnormalities (e.g., subcortical cysts or tumors) were analyzed in the standardized brain. The frontal (proportion: 27.8% [95%CI: 25.7 to 29.9%]) and temporal lobes (42.5% [95%CI: 40.2 to 44.9%]) had a greater probability of including resected electrode sites compared to the parietal (17.7% [95%CI: 15.9 to 19.5%]) and occipital lobes (12.0% [95%CI: 10.5 to 13.5%]; **Table 1**).

Fifty-two patients underwent both pre and postoperative CELF assessments. One of the 52 patients completed the postoperative evaluation of receptive language function alone due to the time constraint (**Supplementary Table S1**). The mean postoperative changes in CLS, RLI, and ELI standard scores were −0.1 (SD: ±9.8 in 51 patients), −1.4 (SD: ±10.6 in 52 patients), and −0.1 (SD: 9.3 in 51 patients). Fourteen among the 51 patients showed a decline in CLS by more than five standard points.

### Concordance between ESM and iEEG high gamma-based mapping

**Figure 4** demonstrates the spatial concordance between language areas defined by ESM and those by high gamma-based mapping. The probability of ESM-induced receptive aphasia at a given cortical point was highly correlated to the degree of significant auditory naming-related high gamma augmentation during the 600-ms periods after sentence onset (rho: +0.46; p<0.001) and before sentence offset (rho: +0.43; p<0.001). Likewise, the probability of ESM-induced auditory expressive aphasia was correlated to auditory naming-related high gamma augmentation during the 600-ms periods after sentence offset (rho: +0.29; p<0.001) and before response onset (rho: +0.29; p<0.001). The probability of ESM-induced visual expressive aphasia was correlated to picture naming-related high gamma augmentation during the 600-ms period after stimulus onset (rho: +0.13; p<0.001). The probability of ESM-induced speech arrest was highly correlated to auditory naming-related high gamma augmentation during the 600-ms period before response onset (rho: +0.53; p<0.001).

### Multivariate regression models incorporating iEEG high gamma augmentation

Multivariate regression models incorporating auditory naming-related high gamma augmentation predicted the postoperative changes in CLS (r^2^ = 0.37; p = 0.015), RLI (r^2^ = 0.43; p = 0.003), and ELI (r^2^ = 0.32; p = 0.048**; Supplementary Table S2**). Higher ‘maximum resected high gamma’ was independently associated with greater decline in CLS (β = −0.09; t = −3.03; p = 0.004) and ELI (β = −0.08; t = −2.63; p = 0.01), but not in RLI (β = −0.04; t = −1.20; p = 0.24). In other words, each 1% amplitude increase at the resected site showing the largest high gamma response resulted in a more severe postoperative decline in CLS by 0.09.

Multivariate regression models incorporating picture naming-related high gamma augmentation likewise predicted the postoperative changes in RLI (r^2^ = 0.50; p < 0.001), but not in CLS (r^2^ = 0.29; p = 0.109) or ELI (r^2^ = 0.27; p = 0.160; **Supplementary Table S3**). Higher ‘maximum resected high gamma’ was independently associated with greater decline in RLI (β = −0.04; t = −2.25; p = 0.030), but not in CLS (β = −0.04; t = −1.82; p = 0.077) or ELI (β = −0.04; t = −1.67; p=0.103).

### Multivariate regression models incorporating iEEG lower frequency band modulations

Each of the multivariate regression models incorporating naming-related low gamma augmentation, beta attenuation, and alpha attenuation during auditory or picture naming task likewise predicted the postoperative changes in CLS, RLI, and ELI (p < 0.05; r^2^ ranging from 0.32 to 0.50; **Supplementary Tables S4-S9**).

### Machine learning-based prediction of postoperative neuropsychological performance

The boosted tree ensemble model, incorporating the aforementioned 16 auditory naming-related iEEG amplitude variables in addition to the dominant hemisphere variable, predicted patients showing a postoperative >5-point CLS decline with an accuracy of 0.80 and area under the curve of 0.65. As shown in **Fig. 5**, this model can highlight cortical sites predicted to result in a >5-point CLS decline, if resected, on the *<u>individual</u>* surface image. **Figure 6A** visualizes the *<u>group-level</u>* probability of a >5-point CLS decline resulting from the virtual resection of a given cortical point on the FreeSurfer averaged surface image. This group-level atlas suggests that resection of the left hemispheric regions, particularly the posterior portions of the temporal neocortices, would increase the risk of language decline. **Figure 6C** visualizes the relative contribution of the 17 variables mentioned above to the prediction model. Resection of sites showing high gamma augmentation during the 600-ms period after stimulus offset had the most substantial contribution to the improved prediction. The relative risk of language decline related to virtual resection of ESM-defined language sites, compared to that of sites outside, was 5.27 (95%CI: 3.47 to 8.02; **Fig. 7**). Virtual resection of ESM-defined receptive language sites maximally increased the relative risk up to 15.9 (95%CI: 9.6-26.3).

**Figure 6.**
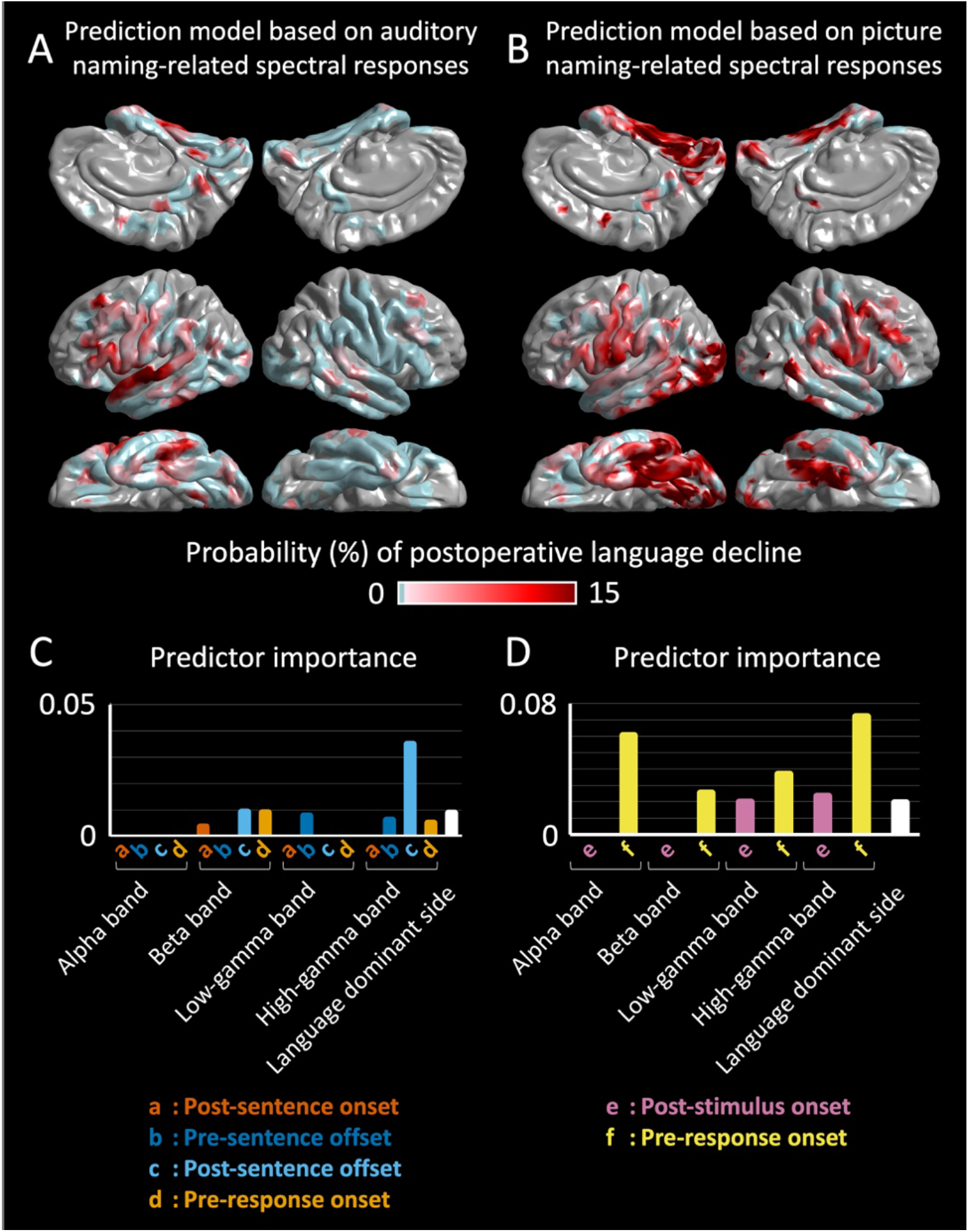
3D brain atlas visualizing the group-level probability of postoperative language decline. The averaged surface image presents the group-level probability of a >5-point CLS decline resulting from the virtual resection of a given cortical point. The group-level probability on each atlas was computed with the boosted tree ensemble model incorporating **A,** auditory naming-related or **B,** picture naming-related iEEG amplitude modulation variables in addition to the dominant hemisphere variable. **C** and **D**, Bar charts visualize the relative contribution of each variable to the prediction model providing atlas **A** and **B**.

**Figure 7.**
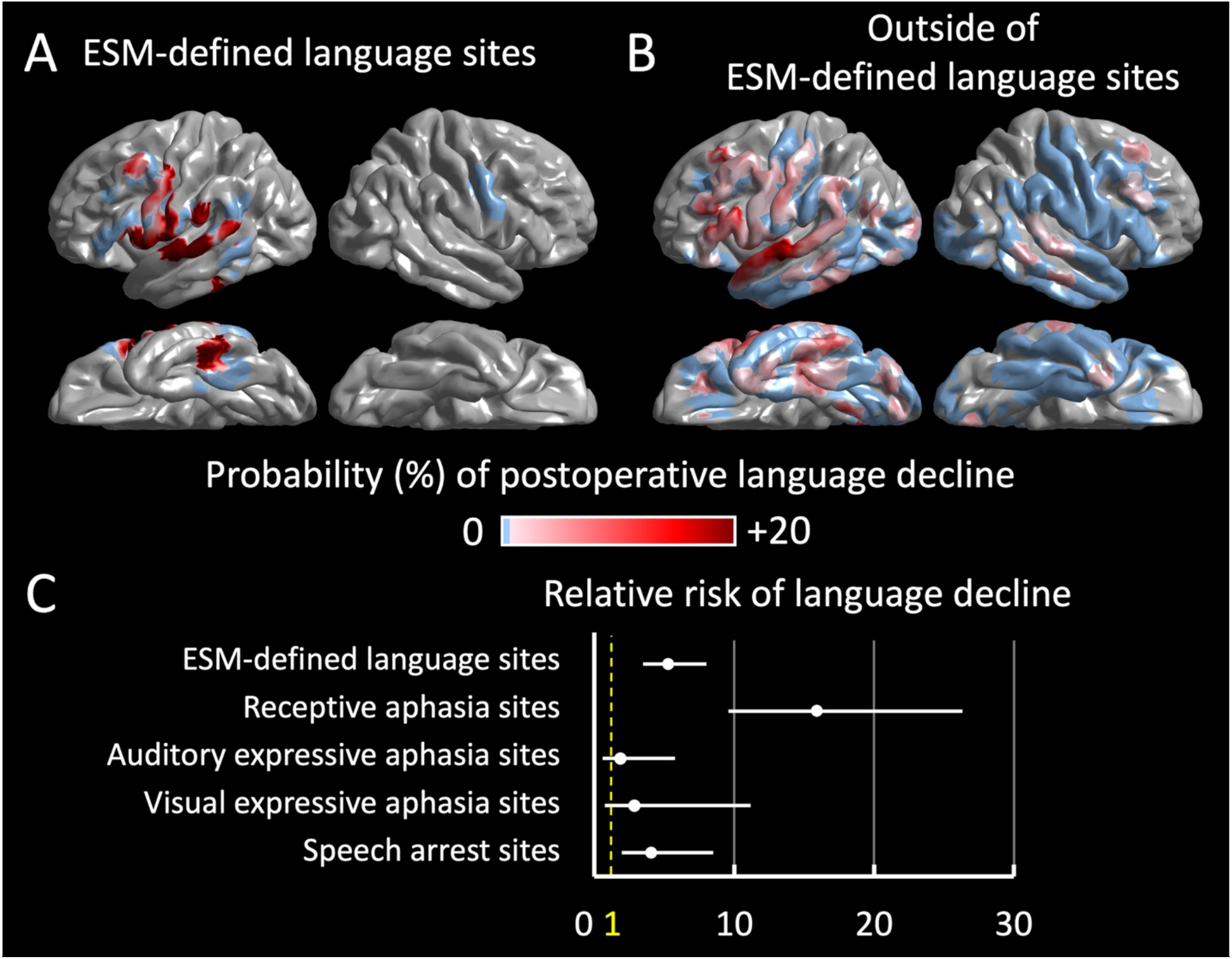
Relative risk of language decline related to the virtual resection of ESM-defined language sites. The boosted tree ensemble model, incorporating auditory naming-related spectral responses, presents the group-level probability of a >5-point CLS decline resulting from *<u>virtual resection</u>* of a given electrode site. **A,** Virtual resection of an ESM-defined language site (i.e., cortical sites with ESM-induced receptive aphasia, auditory expressive aphasia, visual expressive aphasia or speech arrest). **B,** Virtual resection of a site outside the ESM-defined language areas. **C,** The relative risk (95%CI) of language decline related to the virtual resection of an ESM-defined language site, compared to that of a site outside.

The boosted tree ensemble model, incorporating the eight picture naming-related iEEG amplitude variables (**Fig. 6B**) in addition to the dominant hemisphere variable, failed to predict patients showing a >5-point CLS decline with significance (accuracy: 0.73; area under the curve: 0.50).

## DISCUSSION

### Causal significance of our prediction models

This prospective study clarified the causal relationship between resection of sites showing naming-related iEEG amplitude modulations within the language-dominant hemisphere and postoperative changes in objectively-measured neuropsychological performance. We have generated the prediction models that neurosurgeons can utilize before and during the surgical procedures at their own epilepsy centers. The machine learning-based prediction model runs on an ordinary laptop computer with MATLAB installed (**Supplementary .mat file**). All variables incorporated in the prediction models can be determined before the completion of resective surgery. In other words, we did not use measures available only after surgery. It is reasonable to expect that seizure control and reduction of antiepileptic drugs *<u>after surgery</u>* would be associated with improved postoperative language development.^53–56^ However, such postoperative measures can play only a correlative role in characterizing the language function after surgery but cannot be used to predict future symptoms.

### Independent utility of high gamma modulations in predicting neuropsychological outcomes

High gamma-based mapping had a predictive value independent of the effects of each patient’s epilepsy and neuroimaging profiles. Expressly, the regression-based model indicated that greater naming-related high gamma augmentation in the resected dominant hemispheric region accurately predicted a more severe decline in language function after surgery. Our analysis successfully provided novel evidence of the *<u>biological gradient</u>* between naming-related high gamma augmentation and underlying language function. Thereby, the handedness and anatomical MRI lesion determined the language dominant hemisphere (see the Methods Section; ^20,28,29^. The multivariate regression analysis adequately controlled the effects of covariate factors suggested being associated with neuropsychological decline. Investigators have indicated that the risk factors of postoperative language decline include more extensive resection, older age, absence of an MRI-visible lesion, and higher preoperative neuropsychological performance.^33,57–59^ Our multivariate analysis also took into account the time interval between surgery and postoperative neuropsychological assessment (**Table 1**). It is plausible to expect that a given patient recovers and develops language skills as a function of time after surgery. Nonetheless, the present study did not find a significant effect of the time interval on neuropsychological performance. This observation could be attributed to a restriction of range of time post-surgery relative to the typical time course of recovery due to physiologic recovery or reorganization.

Our multivariate regression analysis indicated the dissociative relationship between the amplitude modulations triggered by specific naming tasks and the predicted neuropsychological domain scores. Auditory naming-related high gamma mapping independently predicted the core and expressive language function after surgery, whereas picture naming data did the receptive language function (**Supplementary Tables S2 and S3**). This novel observation can be attributed to the extent of task-related neural activation and the underlying language function. Previous iEEG and functional MRI studies have suggested that the frontal lobe of the language-dominant hemisphere is more extensively and intensively activated by auditory naming task, whereas the ventral temporal-occipital regions are activated by picture naming task.^29,60^ Previous ESM studies have suggested that the left frontal lobe primarily exerts expressive language function and vocalization, whereas the posterior temporal and occipital regions play roles also in the perception and semantic understanding of language stimuli.^3–5,37,61–63^ Our observation does not suggest that preoperative assessment of gamma augmentation with a naming task is superior to the other assessment methods in predicting language outcome.

We have successfully generated the regression- and machine learning-based models predicting postoperative language outcomes *<u>without</u>* relying on the ESM data (**Fig. 6**). Such models are clinically significant because not all patients can have the comprehensive ESM completed for various reasons. Clinicians may not be able to initiate the ESM while patients are prone to develop stimulation-induced seizures due to the reduction or discontinuation of AEDs.^19^ Children may not sustain attentiveness or participation during an hour-long ESM mapping.^8^ Our prediction model based on iEEG high gamma augmentation triggered by naming tasks would be clinically useful because it could partially complement ESM assessment before resective surgery.

In turn, we internally validated our machine learning-based model by assessing the simulated incidence of language impairment resulting from *<u>virtual resection</u>* of ESM-defined language sites (**Fig. 7**). According to the model, resection of an ESM-defined language site, if performed, would have increased the relative risk of a >5-point CLS decline by 5.27. Specifically, virtual resection of ESM-defined receptive language sites maximally increased the relative risk up to 15.90. Our machine learning-based model may provide prognostic information additive to the ESM because it inferred that resection of ESM language-*<u>negative</u>* sites in the left perisylvian regions could still result in a postoperative impairment. A meta-analysis of 15 studies suggests that high gamma mapping may exhibit language areas more extensively than ESM does in pediatric cohorts.^7^ Because we performed ESM using bipolar stimulation, we cannot rule out the possibility that only one of the pair of electrode sites would have been responsible for the symptom elicited during ESM.

### The innovation of our iEEG analysis and prediction model

Our machine learning-based model visualizes the site, which, if removed, would result in a language impairment at the group and individual levels. The group-level 3D atlas (**Fig 6**) visualizes the probability of a postoperative language decline resulting from resection of given cortical sites. The atlas can be readily utilized for counseling and education of patients, students, and healthcare providers. The individual-level model (**Fig 5**) can be used to simulate the postoperative language outcome resulting from the planned resection for a given patient. The accuracy of predicting patients developing a >5-point CLS decline was 0.80 after five-fold crossvalidation. Additional large and diverse datasets will provide an outstanding opportunity to externally validate our prediction models.^64^ At several tertiary epilepsy centers, we currently collect the iEEG, MRI, and neuropsychological datasets, characterized by different electrode types (e.g., depth electrodes), iEEG sampling approach (e.g., more restricted spatial sampling), age group (e.g., adult-dominant cohorts), and spoken language (i.e., other than English).^65^

In the present study, our innovative analysis systematically excluded the time-frequency bins affected by interictal spike discharges, which would randomly take place without being time-locked to stimuli or responses (**Fig. 3**). This method effectively minimized the observation of high gamma augmentation not attributed to task-related neural activations.^28^ Interictal spike discharges are accompanied by a temporary boost of broadband amplitude, including a 30-85 Hz band.^42,43^ Thus, interictal spike discharges, if not removed from the analysis, may undesirably inflate the high gamma amplitudes at non-eloquent cortices, particularly within the SOZ.^66^ We are pleased to share our MATLAB code with investigators who want to replicate our analytic method.

### Methodological considerations

The present study excluded the direct effect of interictal spike discharges on the measurement of naming-related high gamma augmentation at each electrode site. Still, our timefrequency analysis may not consider the indirect impacts of slow-wave accompanying a given spike. Previous iEEG studies reported that slow-wave discharges immediately following spike discharges appeared to reduce task-related high gamma activation in trials affected by spike-and-slow wave discharges.^67,68^ The optimal criteria for exclusion of the effects of epileptiform discharges remain to be determined.

iEEG recording inevitably suffers from a sampling limitation. Thus, we are aware of the possibility that an unsampled cortical site may have generated the true maximum spectral responses. Investigators have looked for noninvasive neurophysiological biomarkers to localize the language areas throughout the cortical convexity. However, the unavoidable occurrence of electromyographic artifacts originating from ocular and temporal muscles during spontaneous saccades and overt responses make the noninvasive high gamma-based language mapping challenging.^69,70^ Our multivariate regression analysis indicated the utility of naming-related alpha/beta attenuation in predicting postoperative language outcomes (**Supplementary Tables S6-S9**). Our study supports the potential role of task-related alpha/beta attenuation measured noninvasively in presurgical evaluation.^71^

The prediction performance of our machine learning-based model inevitably depends on the characteristics of the training data and the strength of model fitness to those data. A large proportion of our study patients had cortical resection involving the frontal or temporal lobe. Thus, our group-level atlas (**Fig. 6**) is expected to provide a more reliable prediction for patients with frontal or temporal lobe epilepsy than those with parietal or occipital lobe epilepsy. It is not reasonable to expect that a single diagnostic test would have a very high diagnostic accuracy approaching 100% in predicting the postoperative language function. Seizure control after surgery is associated with better cognitive development,^33,57,58,72^ but clinicians do not have the seizure outcome before surgery. Thus, further studies also incorporating epilepsy biomarkers capable of predicting postoperative seizure outcome are warranted to improve our prediction model. The promising iEEG candidate biomarkers include high-frequency oscillations (HFOs)^73,74^ and crossfrequency coupling between HFOs and slow waves.^40,75^

The present study did not include patients at the age of four years or younger. We expect that substantial proportions of such young patients will fail to complete an overt naming task satisfactorily. Thus, one would need to establish the language mapping without relying on the child’s attentive participation. Investigators have successfully recorded task-free high gamma augmentation associated with spontaneous cooing and babbling^76^ and during passive listening.^7,77^ Measurement of spectral responses to single-pulse electrical stimulation also has the potential to localize the network supporting speech and language^78–80^ Additional measures are expected to improve the accuracy of machine learning-based prediction models.

## Supporting information

Supplementary material .mat format

## ACKNOWLEDGMENTS

We are grateful to Karin Halsey, BS, REEGT. and Jamie MacDougall, RN, BSN, CPN at Children’s Hospital of Michigan for the collaboration and assistance in performing the studies described above.

## Author contributions

Dr. Asano has full access to all the data in the study and takes responsibility for the integrity of the data and the accuracy of the data analysis.

Concept and design: Sonoda, Asano

Acquisition of data: Rothermel, Carlson, Luat, Sood, Asano.

Analysis, or interpretation of data: Sonoda, Rothermel, Jeong, Lee, Hayashi, Asano.

Drafting of the manuscript: Sonoda, Rothermel, Asano.

Critical revision of the manuscript for important intellectual content: Sonoda, Rothermel, Asano.

Statistical analysis: Sonoda, Asano.

Machine learning: Sonoda, Asano.

Obtained funding: Jeong, Asano.

Administrative, technical, or material support: Rothermel, Luat, Sood, Asano.

Supervision: Asano.

## Abbreviations

AEDs: antiepileptic drugs
CELF-4: Clinical Evaluation of Language Fundamentals-Fourth Edition
CI: confidence interval
CLS: Core Language Score
ELI: Expressive Language Index
ESM: electrical stimulation mapping
HGA: high gamma augmentation
iEEG: intracranial electroencephalography
MRI: magnetic resonance imaging
RLI: Receptive Language Index
SD: standard deviation

