## Supplemental document for "Naming-related spectral responses predict neuropsychological outcome after epilepsy surgery"

**Supplementary Figure S1: Age distribution.**

**Supplementary Table S1: Cognitive profiles.**

**Supplementary Table S2: Multivariate regression analysis incorporating *auditory* naming-related *high gamma augmentation*.**

**Supplementary Table S3: Multivariate regression analysis incorporating *picture* naming-related *high gamma augmentation*.**

**Supplementary Table S4: Multivariate regression analysis incorporating *auditory* naming-related *low gamma augmentation*.**

**Supplementary Table S5: Multivariate regression analysis incorporating *picture* naming-related *low gamma augmentation*.**

**Supplementary Table S6: Multivariate regression analysis incorporating *auditory* naming-related *beta attenuation*.**

**Supplementary Table S7: Multivariate regression analysis incorporating *picture* naming-related *beta attenuation*.**

**Supplementary Table S8: Multivariate regression analysis incorporating *auditory* naming-related *alpha attenuation*.**

**Supplementary Table S9: Multivariate regression analysis incorporating *picture* naming-related *alpha attenuation*.**

**Supplementary .mat file:** **The boosted tree ensemble model incorporating auditory naming-related spectral responses for predicting a postoperative decline of language function.**

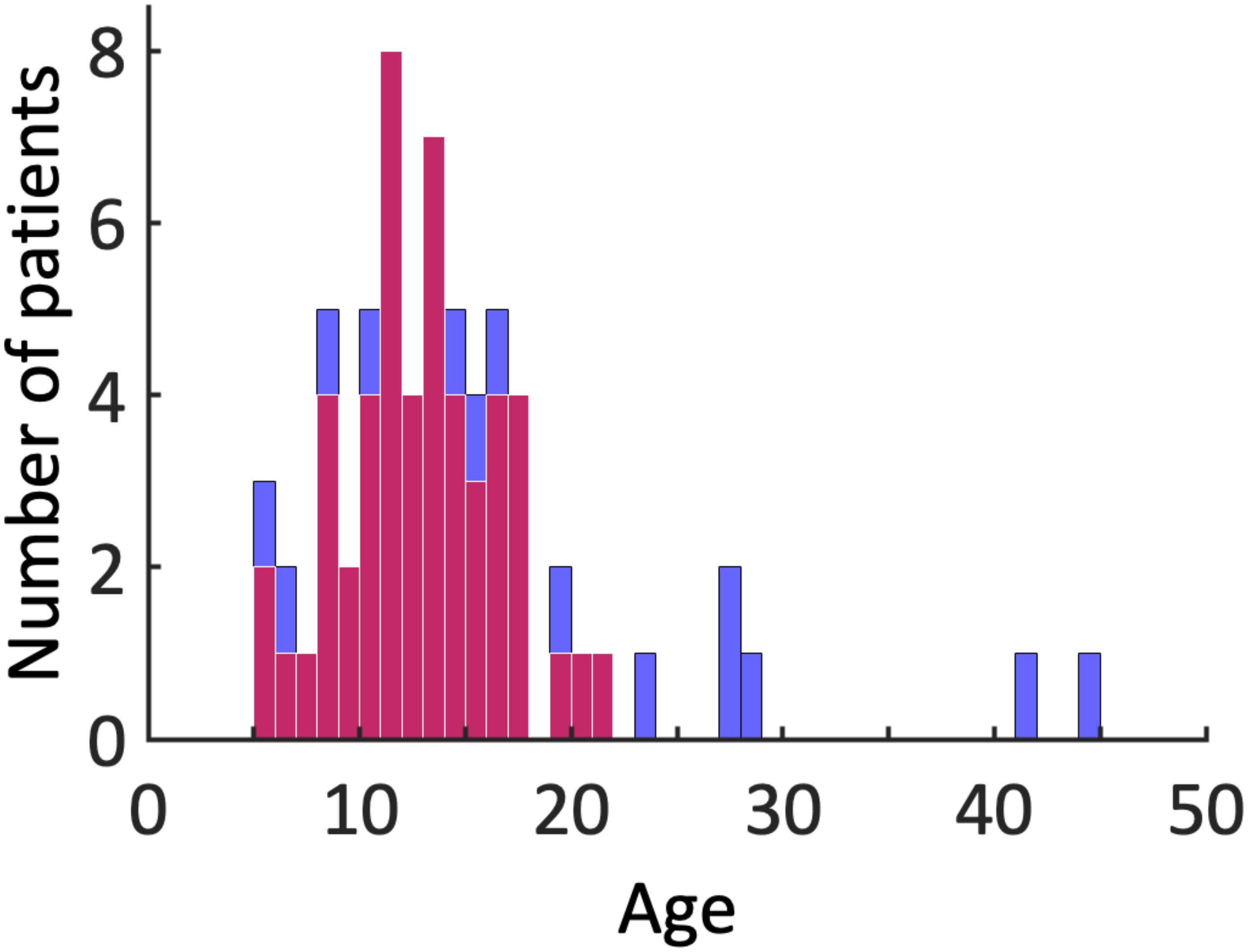


**Supplementary Figure S1: Age histogram.** The histogram presents the distribution of all 65 patients' age included in the present study. Red bars indicate patients whose postoperative decline in Core Language Score (CLS) in CELF-4 could be calculated based on the normative mean and standard deviation;[1] the patients' age followed a normal distribution (Shapiro-Wilk $W$: 0.992; p-value: 0.979; degree of freedom: 51). Blue bars reflect the number of patients in whom postoperative changes in CLS could not be computed due to the lack of pre- or post-operative score.

**Supplementary Table S1: Age distribution.**

| CELF-4 | | n | Mean | SD | Median | IQR |
|---|---|---|---|---|---|---|
| CLS | preoperative | 53 | 81.1 | 22.9 | 79 | 65 - 97.5 |
| | postoperative | 57 | 80.5 | 24.0 | 84 | 65.5 - 99 |
| | Postoperative change | 51 | -0.1 | 9.8 | 0 | -6 - 5.8 |
| RLI | preoperative | 53 | 77.6 | 20.5 | 80 | 58.8 - 90 |
| | postoperative | 58 | 75.7 | 19.4 | 76 | 61 - 90 |
| | Postoperative change | 52 | -1.4 | 10.6 | -1.5 | -6 - 5.5 |
| ELI | preoperative | 53 | 82.7 | 22.1 | 85 | 64.3 - 98.3 |
| | postoperative | 57 | 82.6 | 23.0 | 87 | 65 - 99.5 |
| | Postoperative change | 51 | -0.1 | 9.3 | 0 | -6.8 - 5 |

CELF-4: Clinical Evaluation of Language Fundamentals-Fourth Edition.

CLS: Core Language Score on CELF-4.

ELI: Expressive Language Index on CELF-4.

IQR: interquartile range.

RLI: Receptive Language Index on CELF-4.

SD: standard deviation.

**Supplementary Table S2: Multivariate regression analysis incorporating *auditory* naming-related *high gamma augmentation.***

| Dependent variable | Independent variables | Auditory naming task | | | | |
|---|---|---|---|---|---|---|
| | | β | 95%CI LL | 95%CI UL | *t* | p |
| Postoperative change of CLS<br>$r^2 = 0.37$<br>df = 41<br>p = 0.015 | Maximum resected high gamma | -0.09 | -0.15 | -0.03 | -3.03 | 0.004 |
| | Resection size of language-dominant hemispheric cortex | 0.02 | -0.23 | 0.27 | 0.16 | 0.876 |
| | Age at surgery (years) | 0.77 | 0.04 | 1.50 | 2.13 | 0.039 |
| | Sex (1, if female) | -4.30 | -9.39 | 0.78 | -1.71 | 0.095 |
| | Interval between surgery and postoperative assessment | 0.07 | -1.50 | 1.65 | 0.09 | 0.927 |
| | Number of oral antiepileptic drugs taken preoperatively | 1.91 | -1.42 | 5.24 | 1.16 | 0.254 |
| | MRI-visible cortical lesion (1: if exist) | 2.91 | -2.42 | 8.24 | 1.10 | 0.276 |
| | SOZ location (1: if frontal or temporal) | 4.76 | -1.06 | 10.58 | 1.65 | 0.106 |
| | Preoperative CLS | -0.08 | -0.20 | 0.04 | -1.41 | 0.165 |
| | Constant | -6.49 | -24.35 | 11.37 | -0.73 | 0.467 |
| Postoperative change of RLI<br>$r^2 = 0.43$<br>df = 41<br>p = 0.003 | Maximum resected high gamma | -0.04 | -0.10 | 0.02 | -1.20 | 0.235 |
| | Resection size of language-dominant hemispheric cortex | -0.19 | -0.44 | 0.07 | -1.49 | 0.145 |
| | Age at surgery | 0.38 | -0.36 | 1.12 | 1.04 | 0.304 |
| | Sex (1, if female) | -5.20 | -10.40 | 0.01 | -2.02 | 0.050 |
| | Interval between surgery and postoperative assessment | -1.36 | -2.94 | 0.22 | -1.74 | 0.090 |
| | Number of oral antiepileptic drugs taken preoperatively | 2.08 | -1.30 | 5.46 | 1.24 | 0.220 |
| | MRI-visible cortical lesion (1: if exist) | 2.20 | -3.20 | 7.61 | 0.82 | 0.415 |
| | SOZ location (1: if frontal or temporal) | 6.17 | 0.24 | 12.09 | 2.10 | 0.042 |
| | Preoperative RLI | -0.24 | -0.36 | -0.11 | -3.73 | 0.001 |
| | Constant | 10.48 | -7.06 | 28.02 | 1.21 | 0.235 |
| Postoperative change of ELI<br>$r^2 = 0.31$<br>df = 41<br>p = 0.048 | Maximum resected high gamma | -0.08 | -0.13 | -0.02 | -2.63 | 0.012 |
| | Resection size of language-dominant hemispheric cortex | 0.07 | -0.17 | 0.32 | 0.61 | 0.548 |
| | Age at surgery | 0.70 | -0.01 | 1.41 | 1.98 | 0.054 |
| | Sex (1, if female) | -3.66 | -8.68 | 1.37 | -1.47 | 0.149 |
| | Interval between surgery and postoperative assessment | 0.66 | -0.90 | 2.23 | 0.85 | 0.398 |
| | Number of oral antiepileptic drugs taken preoperatively | 0.81 | -2.48 | 4.10 | 0.50 | 0.621 |
| | MRI-visible cortical lesion (1: if exist) | 3.38 | -1.88 | 8.64 | 1.30 | 0.202 |
| | SOZ location (1: if frontal or temporal) | 5.05 | -0.73 | 10.83 | 1.77 | 0.085 |
| | Preoperative ELI | -0.06 | -0.18 | 0.06 | -0.98 | 0.332 |
| | Constant | -8.01 | -26.38 | 10.35 | -0.88 | 0.383 |

β: estimated regression coefficient. CELF-4: Clinical Evaluation of Language Fundamentals-Fourth Edition. CI: confidence interval. CLS: Core Language Score on CELF-4. df = degree of freedom. ELI: Expressive Language Index on CELF-4. LL: lower limit. MRI, magnetic resonance imaging. p: p value. $r^2$: coefficient of determination. RLI : Receptive Language Index on CELF-4. SOZ: seizure onset zone. *t*: t statistics. UL: upper limit.

**Supplementary Table S3: Multivariate regression analysis incorporating *picture* naming-related *high gamma augmentation.***

| Dependent variable | Independent variables | Picture naming task | | | | |
|---|---|---|---|---|---|---|
| | | β | 95%CI LL | 95%CI UL | *t* | p |
| Postoperative change of CLS<br>$r^2 = 0.29$<br>df = 39<br>p = 0.109 | Maximum resected high gamma | -0.04 | -0.09 | 0.005 | -1.82 | 0.077 |
| | Resection size of language-dominant hemispheric cortex | -0.10 | -0.34 | 0.15 | -0.80 | 0.431 |
| | Age at surgery | 0.93 | 0.09 | 1.76 | 2.25 | 0.030 |
| | Sex (1, if female) | -4.45 | -10.06 | 1.16 | -1.60 | 0.117 |
| | Interval between surgery and postoperative assessment | 0.15 | -1.68 | 1.97 | 0.16 | 0.872 |
| | Number of oral antiepileptic drugs taken preoperatively | 1.95 | -1.96 | 5.85 | 1.01 | 0.319 |
| | MRI-visible cortical lesion (1: if exist) | 2.60 | -3.38 | 8.58 | 0.88 | 0.385 |
| | SOZ location (1: if frontal or temporal) | 2.85 | -3.39 | 9.08 | 0.92 | 0.361 |
| | Preoperative CLS | -0.10 | -0.23 | 0.04 | -1.44 | 0.159 |
| | Constant | -2.76 | -23.24 | 17.72 | -0.27 | 0.787 |
| Postoperative change of RLI<br>$r^2 = 0.50$<br>df = 40<br>p < 0.001 | Maximum resected high gamma | -0.04 | -0.08 | 0.005 | -2.25 | 0.030 |
| | Resection size of language-dominant hemispheric cortex | -0.11 | -0.32 | 0.10 | -1.02 | 0.314 |
| | Age at surgery | 0.59 | -0.15 | 1.32 | 1.62 | 0.113 |
| | Sex (1, if female) | -4.79 | -9.79 | 0.20 | -1.94 | 0.059 |
| | Interval between surgery and postoperative assessment | -0.92 | -2.00 | 0.15 | -1.74 | 0.090 |
| | Number of oral antiepileptic drugs taken preoperatively | 0.89 | -2.46 | 4.24 | 0.53 | 0.596 |
| | MRI-visible cortical lesion (1: if exist) | 1.40 | -3.85 | 6.65 | 0.54 | 0.593 |
| | SOZ location (1: if frontal or temporal) | 5.48 | -0.04 | 11.01 | 2.01 | 0.052 |
| | Preoperative RLI | -0.23 | -0.35 | -0.11 | -3.85 | 0.000 |
| | Constant | 15.67 | -1.61 | 32.96 | 1.83 | 0.074 |
| Postoperative change of ELI<br>$r^2 = 0.27$<br>df = 39<br>p = 0.160 | Maximum resected high gamma | -0.04 | -0.08 | 0.01 | -1.67 | 0.103 |
| | Resection size of language-dominant hemispheric cortex | -0.02 | -0.26 | 0.22 | -0.19 | 0.847 |
| | Age at surgery | 0.86 | 0.06 | 1.66 | 2.18 | 0.035 |
| | Sex (1, if female) | -4.04 | -9.48 | 1.39 | -1.51 | 0.140 |
| | Interval between surgery and postoperative assessment | 0.77 | -1.01 | 2.54 | 0.88 | 0.387 |
| | Number of oral antiepileptic drugs taken preoperatively | 1.17 | -2.61 | 4.96 | 0.63 | 0.534 |
| | MRI-visible cortical lesion (1: if exist) | 3.04 | -2.75 | 8.83 | 1.06 | 0.295 |
| | SOZ location (1: if frontal or temporal) | 3.35 | -2.71 | 9.42 | 1.12 | 0.270 |
| | Preoperative ELI | -0.05 | -0.19 | 0.08 | -0.78 | 0.440 |
| | Constant | -7.08 | -27.84 | 13.67 | -0.69 | 0.494 |

β: estimated regression coefficient. CELF-4: Clinical Evaluation of Language Fundamentals-Fourth Edition. CI: confidence interval. CLS: Core Language Score on CELF-4. df = degree of freedom. ELI: Expressive Language Index on CELF-4. LL: lower limit. MRI, magnetic resonance imaging. p: p value. $r^2$: coefficient of determination. RLI : Receptive Language Index on CELF-4. SOZ: seizure onset zone. *t*: t statistics. UL: upper limit.

**Supplementary Table S4: Multivariate regression analysis incorporating *auditory* naming-related *low gamma augmentation.***

| Dependent variable | Independent variables | Auditory naming task β | 95%CI LL | 95%CI UL | *t* | p |
|---|---|---|---|---|---|---|
| Postoperative change of CLS; $r^2 = 0.39$; df = 41; p = 0.009 | Maximum resected low gamma | -0.09 | -0.15 | -0.04 | -3.30 | 0.002 |
| | Resection size of language-dominant hemispheric cortex | -0.03 | -0.26 | 0.19 | -0.30 | 0.765 |
| | Age at surgery | 0.73 | 0.01 | 1.44 | 2.05 | 0.047 |
| | Sex (1, if female) | -4.22 | -9.22 | 0.78 | -1.70 | 0.096 |
| | Interval between surgery and postoperative assessment | 0.06 | -1.48 | 1.61 | 0.08 | 0.935 |
| | Number of oral antiepileptic drugs taken preoperatively | 2.10 | -1.17 | 5.37 | 1.30 | 0.201 |
| | MRI-visible cortical lesion (1: if exist) | 3.09 | -2.16 | 8.34 | 1.19 | 0.241 |
| | SOZ location (1: if frontal or temporal) | 4.83 | -0.89 | 10.54 | 1.71 | 0.095 |
| | Preoperative CLS | -0.08 | -0.20 | 0.04 | -1.41 | 0.166 |
| | Constant | -6.42 | -23.98 | 11.14 | -0.74 | 0.465 |
| Postoperative change of RLI; $r^2 = 0.43$; df = 41; p = 0.002 | Maximum resected low gamma | -0.04 | -0.10 | 0.02 | -1.43 | 0.161 |
| | Resection size of language-dominant hemispheric cortex | -0.20 | -0.43 | 0.03 | -1.78 | 0.083 |
| | Age at surgery | 0.36 | -0.37 | 1.09 | 1.01 | 0.320 |
| | Sex (1, if female) | -5.15 | -10.32 | 0.02 | -2.01 | 0.051 |
| | Interval between surgery and postoperative assessment | -1.39 | -2.95 | 0.18 | -1.79 | 0.081 |
| | Number of oral antiepileptic drugs taken preoperatively | 2.15 | -1.20 | 5.49 | 1.30 | 0.202 |
| | MRI-visible cortical lesion (1: if exist) | 2.29 | -3.08 | 7.67 | 0.86 | 0.394 |
| | SOZ location (1: if frontal or temporal) | 6.28 | 0.41 | 12.15 | 2.16 | 0.037 |
| | Preoperative RLI | -0.23 | -0.36 | -0.11 | -3.72 | 0.001 |
| | Constant | 10.56 | -6.86 | 27.98 | 1.22 | 0.228 |
| Postoperative change of ELI; $r^2 = 0.34$; df = 41; p = 0.030 | Maximum resected low gamma | -0.08 | -0.14 | -0.02 | -2.91 | 0.006 |
| | Resection size of language-dominant hemispheric cortex | 0.03 | -0.19 | 0.25 | 0.29 | 0.775 |
| | Age at surgery | 0.66 | -0.04 | 1.36 | 1.91 | 0.063 |
| | Sex (1, if female) | -3.58 | -8.52 | 1.37 | -1.46 | 0.152 |
| | Interval between surgery and postoperative assessment | 0.65 | -0.89 | 2.18 | 0.85 | 0.401 |
| | Number of oral antiepileptic drugs taken preoperatively | 0.97 | -2.26 | 4.20 | 0.61 | 0.547 |
| | MRI-visible cortical lesion (1: if exist) | 3.54 | -1.64 | 8.72 | 1.38 | 0.175 |
| | SOZ location (1: if frontal or temporal) | 5.15 | -0.53 | 10.82 | 1.83 | 0.074 |
| | Preoperative ELI | -0.06 | -0.18 | 0.06 | -0.97 | 0.336 |
| | Constant | -7.92 | -25.99 | 10.15 | -0.89 | 0.381 |

β: estimated regression coefficient. CELF-4: Clinical Evaluation of Language Fundamentals-Fourth Edition. CI: confidence interval. CLS: Core Language Score on CELF-4. df = degree of freedom. ELI: Expressive Language Index on CELF-4. LL: lower limit. MRI, magnetic resonance imaging. p: p value. $r^2$: coefficient of determination. RLI : Receptive Language Index on CELF-4. SOZ: seizure onset zone. *t*: t statistics. UL: upper limit.

**Supplementary Table S5: Multivariate regression analysis incorporating *picture* naming-related *low gamma augmentation.***

| Dependent variable | Independent variables | Picture naming task | | | | |
|---|---|---|---|---|---|---|
| | | β | 95%CI LL | 95%CI UL | *t* | p |
| Postoperative change of CLS $r^2 = 0.30$ df = 39 p = 0.091 | Maximum resected low gamma | -0.04 | -0.09 | 0.00 | -1.97 | 0.056 |
| | Resection size of language-dominant hemispheric cortex | -0.10 | -0.34 | 0.14 | -0.84 | 0.408 |
| | Age at surgery | 0.93 | 0.11 | 1.75 | 2.28 | 0.028 |
| | Sex (1, if female) | -4.01 | -9.64 | 1.61 | -1.44 | 0.157 |
| | Interval between surgery and postoperative assessment | 0.20 | -1.59 | 1.98 | 0.22 | 0.825 |
| | Number of oral antiepileptic drugs taken preoperatively | 1.66 | -2.23 | 5.55 | 0.86 | 0.393 |
| | MRI-visible cortical lesion (1: if exist) | 2.98 | -3.00 | 8.96 | 1.01 | 0.319 |
| | SOZ location (1: if frontal or temporal) | 2.59 | -3.58 | 8.77 | 0.85 | 0.401 |
| | Preoperative CLS | -0.10 | -0.23 | 0.04 | -1.47 | 0.148 |
| | Constant | -2.82 | -23.10 | 17.45 | -0.28 | 0.780 |
| Postoperative change of RLI $r^2 = 0.45$ df = 40 p = 0.002 | Maximum resected low gamma | -0.02 | -0.06 | 0.02 | -0.88 | 0.386 |
| | Resection size of language-dominant hemispheric cortex | -0.18 | -0.40 | 0.04 | -1.66 | 0.105 |
| | Age at surgery | 0.44 | -0.33 | 1.20 | 1.16 | 0.253 |
| | Sex (1, if female) | -5.02 | -10.33 | 0.29 | -1.91 | 0.063 |
| | Interval between surgery and postoperative assessment | -0.71 | -1.82 | 0.40 | -1.30 | 0.202 |
| | Number of oral antiepileptic drugs taken preoperatively | 0.89 | -2.66 | 4.44 | 0.51 | 0.615 |
| | MRI-visible cortical lesion (1: if exist) | 1.38 | -4.17 | 6.92 | 0.50 | 0.619 |
| | SOZ location (1: if frontal or temporal) | 5.03 | -0.75 | 10.82 | 1.76 | 0.086 |
| | Preoperative RLI | -0.24 | -0.37 | -0.12 | -3.87 | 0.000 |
| | Constant | 13.53 | -4.59 | 31.65 | 1.51 | 0.139 |
| Postoperative change of ELI $r^2 = 0.27$ df = 39 p = 0.146 | Maximum resected low gamma | -0.04 | -0.08 | 0.01 | -1.77 | 0.085 |
| | Resection size of language-dominant hemispheric cortex | -0.03 | -0.26 | 0.21 | -0.24 | 0.814 |
| | Age at surgery | 0.86 | 0.07 | 1.65 | 2.19 | 0.035 |
| | Sex (1, if female) | -3.68 | -9.14 | 1.79 | -1.36 | 0.182 |
| | Interval between surgery and postoperative assessment | 0.82 | -0.92 | 2.56 | 0.95 | 0.346 |
| | Number of oral antiepileptic drugs taken preoperatively | 0.92 | -2.86 | 4.70 | 0.49 | 0.625 |
| | MRI-visible cortical lesion (1: if exist) | 3.36 | -2.44 | 9.17 | 1.17 | 0.248 |
| | SOZ location (1: if frontal or temporal) | 3.12 | -2.91 | 9.15 | 1.05 | 0.302 |
| | Preoperative ELI | -0.06 | -0.19 | 0.08 | -0.82 | 0.418 |
| | Constant | -7.16 | -27.79 | 13.48 | -0.70 | 0.487 |

β: estimated regression coefficient. CELF-4: Clinical Evaluation of Language Fundamentals-Fourth Edition. CI: confidence interval. CLS: Core Language Score on CELF-4. df = degree of freedom. ELI: Expressive Language Index on CELF-4. LL: lower limit. MRI, magnetic resonance imaging. p: p value. $r^2$: coefficient of determination. RLI : Receptive Language Index on CELF-4. SOZ: seizure onset zone. *t*: t statistics. UL: upper limit.

**Supplementary Table S6: Multivariate regression analysis incorporating *auditory* naming-related *beta attenuation.***

| Dependent variable | Independent variables | Auditory naming task | | | | |
|---|---|---|---|---|---|---|
| | | β | 95%CI | | *t* | p |
| | | | LL | UL | | |
| Postoperative change of CLS<br><br>$r^2 = 0.38$<br>df = 41<br>p = 0.012 | Maximum resected beta | -0.09 | -0.14 | -0.03 | -3.17 | 0.003 |
| | Resection size of language-dominant hemispheric cortex | -0.03 | -0.26 | 0.20 | -0.23 | 0.817 |
| | Age at surgery | 0.79 | 0.06 | 1.51 | 2.19 | 0.034 |
| | Sex (1, if female) | -4.15 | -9.20 | 0.90 | -1.66 | 0.105 |
| | Interval between surgery and postoperative assessment | 0.10 | -1.45 | 1.66 | 0.13 | 0.896 |
| | Number of oral antiepileptic drugs taken preoperatively | 2.06 | -1.24 | 5.35 | 1.26 | 0.214 |
| | MRI-visible cortical lesion (1: if exist) | 3.07 | -2.22 | 8.36 | 1.17 | 0.248 |
| | SOZ location (1: if frontal or temporal) | 4.76 | -1.00 | 10.52 | 1.67 | 0.103 |
| | Preoperative CLS | -0.08 | -0.20 | 0.04 | -1.37 | 0.179 |
| | Constant | -7.42 | -25.11 | 10.27 | -0.85 | 0.402 |
| Postoperative change of RLI<br><br>$r^2 = 0.43$<br>df = 41<br>p = 0.003 | Maximum resected beta | -0.04 | -0.09 | 0.02 | -1.27 | 0.210 |
| | Resection size of language-dominant hemispheric cortex | -0.20 | -0.44 | 0.03 | -1.77 | 0.085 |
| | Age at surgery | 0.39 | -0.35 | 1.12 | 1.06 | 0.295 |
| | Sex (1, if female) | -5.14 | -10.34 | 0.06 | -2.00 | 0.053 |
| | Interval between surgery and postoperative assessment | -1.35 | -2.93 | 0.22 | -1.74 | 0.089 |
| | Number of oral antiepileptic drugs taken preoperatively | 2.14 | -1.23 | 5.50 | 1.28 | 0.206 |
| | MRI-visible cortical lesion (1: if exist) | 2.27 | -3.13 | 7.66 | 0.85 | 0.402 |
| | SOZ location (1: if frontal or temporal) | 6.18 | 0.28 | 12.08 | 2.12 | 0.040 |
| | Preoperative RLI | -0.23 | -0.36 | -0.11 | -3.70 | 0.001 |
| | Constant | 10.09 | -7.38 | 27.57 | 1.17 | 0.250 |
| Postoperative change of ELI<br>$r^2 = 0.33$<br>df = 41<br>p = 0.041 | Maximum resected beta | -0.07 | -0.13 | -0.02 | -2.73 | 0.009 |
| | Resection size of language-dominant hemispheric cortex | 0.03 | -0.19 | 0.26 | 0.30 | 0.762 |
| | Age at surgery | 0.71 | 0.00 | 1.42 | 2.03 | 0.049 |
| | Sex (1, if female) | -3.53 | -8.53 | 1.47 | -1.43 | 0.162 |
| | Interval between surgery and postoperative assessment | 0.69 | -0.86 | 2.24 | 0.90 | 0.375 |
| | Number of oral antiepileptic drugs taken preoperatively | 0.94 | -2.33 | 4.20 | 0.58 | 0.565 |
| | MRI-visible cortical lesion (1: if exist) | 3.51 | -1.73 | 8.74 | 1.35 | 0.184 |
| | SOZ location (1: if frontal or temporal) | 5.05 | -0.68 | 10.79 | 1.78 | 0.083 |
| | Preoperative ELI | -0.06 | -0.18 | 0.07 | -0.95 | 0.347 |
| | Constant | -8.77 | -27.02 | 9.47 | -0.97 | 0.337 |

β: estimated regression coefficient. CELF-4: Clinical Evaluation of Language Fundamentals-Fourth Edition. CI: confidence interval. CLS: Core Language Score on CELF-4. df = degree of freedom. ELI: Expressive Language Index on CELF-4. LL: lower limit. MRI, magnetic resonance imaging. p: p value. $r^2$: coefficient of determination. RLI : Receptive Language Index on CELF-4. SOZ: seizure onset zone. *t*: t statistics. UL: upper limit.

**Supplementary Table S7: Multivariate regression analysis incorporating *picture* naming-related *beta attenuation.***

| Dependent variable | Independent variables | Picture naming task | | | | |
|---|---|---|---|---|---|---|
| | | β | 95%CI LL | 95%CI UL | *t* | p |
| Postoperative change of CLS<br>$r^2$ = 0.40<br>df = 39<br>p = 0.011 | Maximum resected beta | -0.09 | -0.15 | -0.04 | -3.30 | 0.002 |
| | Resection size of language-dominant hemispheric cortex | -0.06 | -0.28 | 0.16 | -0.53 | 0.599 |
| | Age at surgery | 0.93 | 0.18 | 1.67 | 2.52 | 0.016 |
| | Sex (1, if female) | -5.03 | -10.17 | 0.11 | -1.98 | 0.055 |
| | Interval between surgery and postoperative assessment | -0.04 | -1.69 | 1.60 | -0.05 | 0.958 |
| | Number of oral antiepileptic drugs taken preoperatively | 3.25 | -0.44 | 6.94 | 1.78 | 0.083 |
| | MRI-visible cortical lesion (1: if exist) | 2.90 | -2.60 | 8.41 | 1.07 | 0.293 |
| | SOZ location (1: if frontal or temporal) | 5.11 | -0.84 | 11.06 | 1.74 | 0.090 |
| | Preoperative CLS | -0.08 | -0.20 | 0.04 | -1.30 | 0.200 |
| | Constant | -1.01 | -19.81 | 17.79 | -0.11 | 0.914 |
| Postoperative change of RLI<br>$r^2$ = 0.47<br>df = 40<br>p = 0.001 | Maximum resected beta | -0.04 | -0.10 | 0.02 | -1.45 | 0.156 |
| | Resection size of language-dominant hemispheric cortex | -0.16 | -0.37 | 0.05 | -1.51 | 0.138 |
| | Age at surgery | 0.44 | -0.29 | 1.18 | 1.22 | 0.231 |
| | Sex (1, if female) | -5.46 | -10.59 | -0.34 | -2.15 | 0.037 |
| | Interval between surgery and postoperative assessment | -0.82 | -1.92 | 0.29 | -1.49 | 0.144 |
| | Number of oral antiepileptic drugs taken preoperatively | 1.54 | -1.98 | 5.05 | 0.88 | 0.382 |
| | MRI-visible cortical lesion (1: if exist) | 1.34 | -4.10 | 6.77 | 0.50 | 0.622 |
| | SOZ location (1: if frontal or temporal) | 6.14 | 0.20 | 12.07 | 2.09 | 0.043 |
| | Preoperative RLI | -0.23 | -0.36 | -0.11 | -3.73 | 0.001 |
| | Constant | 14.32 | -3.51 | 32.15 | 1.62 | 0.112 |
| Postoperative change of ELI<br>$r^2$ = 0.34<br>df = 39<br>p = 0.037 | Maximum resected beta | -0.08 | -0.13 | -0.02 | -2.79 | 0.008 |
| | Resection size of language-dominant hemispheric cortex | 0.00 | -0.22 | 0.23 | 0.04 | 0.970 |
| | Age at surgery | 0.86 | 0.12 | 1.59 | 2.36 | 0.023 |
| | Sex (1, if female) | -4.57 | -9.68 | 0.54 | -1.81 | 0.078 |
| | Interval between surgery and postoperative assessment | 0.63 | -1.01 | 2.28 | 0.78 | 0.442 |
| | Number of oral antiepileptic drugs taken preoperatively | 2.25 | -1.42 | 5.93 | 1.24 | 0.223 |
| | MRI-visible cortical lesion (1: if exist) | 3.25 | -2.22 | 8.72 | 1.20 | 0.236 |
| | SOZ location (1: if frontal or temporal) | 5.25 | -0.70 | 11.20 | 1.78 | 0.082 |
| | Preoperative ELI | -0.04 | -0.17 | 0.09 | -0.68 | 0.500 |
| | Constant | -5.64 | -25.25 | 13.97 | -0.58 | 0.564 |

β: estimated regression coefficient. CELF-4: Clinical Evaluation of Language Fundamentals-Fourth Edition. CI: confidence interval. CLS: Core Language Score on CELF-4. df = degree of freedom. ELI: Expressive Language Index on CELF-4. LL: lower limit. MRI, magnetic resonance imaging. p: p value. $r^2$: coefficient of determination. RLI : Receptive Language Index on CELF-4. SOZ: seizure onset zone. *t*: t statistics. UL: upper limit.

**Supplementary Table S8: Multivariate regression analysis incorporating *auditory* naming-related *alpha attenuation.***

| Dependent variable | Independent variables | Auditory naming task | | | | |
|---|---|---|---|---|---|---|
| | | β | 95%CI LL | 95%CI UL | *t* | p |
| Postoperative change of CLS<br>$r^2 = 0.38$<br>df = 41<br>p = 0.012 | Maximum resected alpha | -0.08 | -0.14 | -0.03 | -3.17 | 0.003 |
| | Resection size of language-dominant hemispheric cortex | -0.03 | -0.26 | 0.20 | -0.24 | 0.809 |
| | Age at surgery | 0.78 | 0.06 | 1.50 | 2.17 | 0.035 |
| | Sex (1, if female) | -4.09 | -9.14 | 0.96 | -1.63 | 0.110 |
| | Interval between surgery and postoperative assessment | 0.09 | -1.46 | 1.65 | 0.12 | 0.904 |
| | Number of oral antiepileptic drugs taken preoperatively | 1.97 | -1.33 | 5.27 | 1.21 | 0.235 |
| | MRI-visible cortical lesion (1: if exist) | 3.08 | -2.22 | 8.37 | 1.17 | 0.247 |
| | SOZ location (1: if frontal or temporal) | 4.82 | -0.95 | 10.59 | 1.69 | 0.099 |
| | Preoperative CLS | -0.08 | -0.20 | 0.04 | -1.38 | 0.174 |
| | Constant | -7.11 | -24.80 | 10.59 | -0.81 | 0.422 |
| Postoperative change of RLI<br>$r^2 = 0.43$<br>df = 41<br>p = 0.003 | Maximum resected alpha | -0.04 | -0.09 | 0.02 | -1.31 | 0.196 |
| | Resection size of language-dominant hemispheric cortex | -0.20 | -0.43 | 0.03 | -1.75 | 0.087 |
| | Age at surgery | 0.39 | -0.35 | 1.12 | 1.06 | 0.296 |
| | Sex (1, if female) | -5.10 | -10.30 | 0.09 | -1.98 | 0.054 |
| | Interval between surgery and postoperative assessment | -1.36 | -2.93 | 0.21 | -1.76 | 0.087 |
| | Number of oral antiepileptic drugs taken preoperatively | 2.10 | -1.27 | 5.46 | 1.26 | 0.215 |
| | MRI-visible cortical lesion (1: if exist) | 2.28 | -3.12 | 7.67 | 0.85 | 0.399 |
| | SOZ location (1: if frontal or temporal) | 6.23 | 0.33 | 12.13 | 2.13 | 0.039 |
| | Preoperative RLI | -0.23 | -0.36 | -0.11 | -3.71 | 0.001 |
| | Constant | 10.26 | -7.21 | 27.72 | 1.19 | 0.242 |
| Postoperative change of ELI<br>$r^2 = 0.33$<br>df = 41<br>p = 0.043 | Maximum resected alpha | -0.07 | -0.12 | -0.02 | -2.70 | 0.010 |
| | Resection size of language-dominant hemispheric cortex | 0.03 | -0.20 | 0.26 | 0.28 | 0.777 |
| | Age at surgery | 0.71 | 0.00 | 1.42 | 2.01 | 0.051 |
| | Sex (1, if female) | -3.48 | -8.50 | 1.53 | -1.40 | 0.168 |
| | Interval between surgery and postoperative assessment | 0.68 | -0.87 | 2.24 | 0.89 | 0.379 |
| | Number of oral antiepileptic drugs taken preoperatively | 0.87 | -2.41 | 4.14 | 0.54 | 0.595 |
| | MRI-visible cortical lesion (1: if exist) | 3.51 | -1.73 | 8.75 | 1.35 | 0.184 |
| | SOZ location (1: if frontal or temporal) | 5.08 | -0.67 | 10.84 | 1.78 | 0.082 |
| | Preoperative ELI | -0.06 | -0.18 | 0.06 | -0.97 | 0.340 |
| | Constant | -8.51 | -26.79 | 9.77 | -0.94 | 0.353 |

β: estimated regression coefficient. CELF-4: Clinical Evaluation of Language Fundamentals-Fourth Edition. CI: confidence interval. CLS: Core Language Score on CELF-4. df = degree of freedom. ELI: Expressive Language Index on CELF-4. LL: lower limit. MRI, magnetic resonance imaging. p: p value. $r^2$: coefficient of determination. RLI : Receptive Language Index on CELF-4. SOZ: seizure onset zone. *t*: t statistics. UL: upper limit.

**Supplementary Table S9: Multivariate regression analysis incorporating *picture* naming-related *alpha attenuation.***

| Dependent variable | Independent variables | Picture naming task β | 95%CI LL | 95%CI UL | *t* | p |
|---|---|---|---|---|---|---|
| Postoperative change of CLS; $r^2 = 0.41$, df = 39, p = 0.008 | Maximum resected alpha | -0.10 | -0.16 | -0.04 | -3.44 | 0.001 |
| | Resection size of language-dominant hemispheric cortex | -0.05 | -0.27 | 0.17 | -0.47 | 0.643 |
| | Age at surgery | 0.91 | 0.17 | 1.64 | 2.49 | 0.017 |
| | Sex (1, if female) | -5.01 | -10.09 | 0.08 | -1.99 | 0.054 |
| | Interval between surgery and postoperative assessment | -0.05 | -1.68 | 1.58 | -0.06 | 0.952 |
| | Number of oral antiepileptic drugs taken preoperatively | 2.79 | -0.82 | 6.39 | 1.56 | 0.126 |
| | MRI-visible cortical lesion (1: if exist) | 2.87 | -2.58 | 8.32 | 1.06 | 0.294 |
| | SOZ location (1: if frontal or temporal) | 5.02 | -0.85 | 10.88 | 1.73 | 0.092 |
| | Preoperative CLS | -0.08 | -0.21 | 0.04 | -1.39 | 0.171 |
| | Constant | 1.42 | -17.47 | 20.31 | 0.15 | 0.880 |
| Postoperative change of RLI; $r^2 = 0.47$, df = 40, p = 0.001 | Maximum resected alpha | -0.04 | -0.10 | 0.01 | -1.52 | 0.135 |
| | Resection size of language-dominant hemispheric cortex | -0.15 | -0.37 | 0.06 | -1.49 | 0.145 |
| | Age at surgery | 0.44 | -0.30 | 1.17 | 1.21 | 0.235 |
| | Sex (1, if female) | -5.46 | -10.57 | -0.34 | -2.16 | 0.037 |
| | Interval between surgery and postoperative assessment | -0.84 | -1.95 | 0.27 | -1.54 | 0.133 |
| | Number of oral antiepileptic drugs taken preoperatively | 1.36 | -2.10 | 4.83 | 0.79 | 0.431 |
| | MRI-visible cortical lesion (1: if exist) | 1.34 | -4.08 | 6.76 | 0.50 | 0.620 |
| | SOZ location (1: if frontal or temporal) | 6.11 | 0.22 | 12.00 | 2.10 | 0.042 |
| | Preoperative RLI | -0.24 | -0.36 | -0.11 | -3.79 | 0.001 |
| | Constant | 15.47 | -2.59 | 33.53 | 1.73 | 0.091 |
| Postoperative change of ELI; $r^2 = 0.36$, df = 39, p = 0.026 | Maximum resected alpha | -0.08 | -0.14 | -0.03 | -3.00 | 0.005 |
| | Resection size of language-dominant hemispheric cortex | 0.01 | -0.21 | 0.23 | 0.13 | 0.901 |
| | Age at surgery | 0.84 | 0.12 | 1.56 | 2.37 | 0.023 |
| | Sex (1, if female) | -4.55 | -9.59 | 0.50 | -1.82 | 0.076 |
| | Interval between surgery and postoperative assessment | 0.61 | -1.01 | 2.23 | 0.76 | 0.452 |
| | Number of oral antiepileptic drugs taken preoperatively | 1.89 | -1.69 | 5.46 | 1.07 | 0.292 |
| | MRI-visible cortical lesion (1: if exist) | 3.24 | -2.15 | 8.64 | 1.22 | 0.231 |
| | SOZ location (1: if frontal or temporal) | 5.23 | -0.62 | 11.07 | 1.81 | 0.078 |
| | Preoperative ELI | -0.05 | -0.17 | 0.08 | -0.74 | 0.465 |
| | Constant | -3.40 | -23.00 | 16.20 | -0.35 | 0.727 |

β: estimated regression coefficient. CELF-4: Clinical Evaluation of Language Fundamentals-Fourth Edition. CI: confidence interval. CLS: Core Language Score on CELF-4. df = degree of freedom. ELI: Expressive Language Index on CELF-4. LL: lower limit. MRI, magnetic resonance imaging. p: p value. $r^2$: coefficient of determination. RLI : Receptive Language Index on CELF-4. SOZ: seizure onset zone. *t*: t statistics. UL: upper limit.

**Supplementary .mat file: The boosted tree ensemble model incorporating auditory naming-related spectral responses for predicting a postoperative decline of language function.**

This MATLAB-based .mat file contains [**A**] '*trainedModel*' consisting of our boosted tree ensemble model incorporating auditory naming-related amplitude modulations, [**B**] '*exampleDataset1*', and [**C**] '*exampleDataset2*'. **[B]** This example dataset contains 17 variables, including 16 spectral responses, each averaged across 37 ESM-defined receptive aphasia sites. **[C]** This example dataset contains 17 variables, including 16 spectral responses, each averaged across 6448 sites outside the ESM-defined language sites.

To utilize our boosted tree ensemble-based prediction model for future patients, users will run the following statement within the MATLAB command window:

```
yfit = trainedModel.predictFcn(Dataset)
```

For example, users can validate our '*exampleDataset1*' mentioned above by running the following command:

```
yfit = trainedModel.predictFcn(exampleDataset1)
```

Users will need to enter the electrode's dataset including the 17 variables mentioned above. The model will provide its prediction about whether the resection of the electrode site would result in a >5 points CLS decline ('yfit' of 1: predicted to have a language decline).
